# A hyperbolic equation for modelling non-linear responses of forest tree growth to environmental factors

**DOI:** 10.64898/2026.02.27.708557

**Authors:** Patrick Vallet

## Abstract

The influence of environmental factors on the dynamics of tree species can imply non-linear relationships. Some of them exhibit threshold effects. Hyperbolic functions effectively represent ecological processes that display threshold behaviours. However, the mathematical formulation of the hyperbola is complex, which makes its use challenging and its parameters difficult to interpret.

In this article, we propose an efficient mathematical formulation for the hyperbola, one in which all the parameters are independent and easily interpretable. We also provide an R script and a Python script to facilitate the implementation of this hyperbolic formulation in modelling studies.

We then modelled the influence of edaphic and climatic factors on the growth of 18 forest tree species widely distributed across Europe based on a dataset of 8,330 plots from the French National Forest Inventory. Our hyperbolic function allowed us to identify the threshold effects of summer climatic constraints on forest growth for several species. In particular, we found negative effects for soil water deficit and maximum summer temperature, although for several species these effects only appear beyond a certain level of constraint. Accounting for such threshold effects is crucial to improve our ability to understand and predict forest ecosystem responses in the context of climate change.

## 1. Introduction

Understanding ecological processes often rely on mathematical models that can capture the complexity of living systems. Among these models, hyperbolic functions hold a special place, as they can account for the non-linear behaviour frequently observed in ecological dynamics. The ecological relevance of hyperbolic functions thus lies in their ability to represent a rapid increase followed by an asymptotic slowdown, or conversely, a sharp decline beyond a threshold. In this sense, using hyperbolic functions in ecology extends beyond mathematical formalism; such functions are powerful tools for anticipating tipping points in species’ response to environmental factors.

In forest ecology, hyperbolas have also been used to model the increase in tree basal area as a function of their circumference (Deleuze et al., 2004). In this case, tree growth remains very low up to a certain threshold, beyond which access to light enables an almost linear increase in tree size. This corresponds to a hyperbola with an initial horizontal asymptote, followed by a positive oblique asymptote. Dhôte and de Hercé (1994) also used a hyperbolic model to describe the relationship between tree circumference and height in even-aged stands. Here, the hyperbola first exhibits an oblique asymptote representing the linear relationship between circumference and height, then a horizontal asymptote indicating height saturation beyond a given circumference value. In both cases, the hyperbolic function not only accurately captures the relationship between the variables but also avoids singularities in the equation, which can hinder the convergence of regression algorithms, such as those that may arise with segmented lines.

However, the canonical equation of a hyperbola only describes the special case in which the axes are aligned with the coordinate axes. The formulation of hyperbolas that are not centred at the origin and whose branches are neither horizontal nor vertical involves significant equation complexity, making the parameters difficult to interpret. This complexity limits the intuitiveness of both analysis and visualisation, and interpretability is essential in applied contexts, for example, when studying the links between ecological functions and environmental factors.

In this article, we propose a mathematical formulation of the hyperbola in which the parameters are decorrelated from each other and easily interpretable. We then apply our hyperbolic equation to modelling forest stand growth in France. We hypothesized that forest growth would respond non-linearly to summer climatic constraints: that is, summer constraints would be negligible up to a certain threshold, beyond which they would exert a negative effect. We assumed that the phenomenon could be modelled with a hyperbolic function. Accurately accounting for summer constraints is particularly important in the context of climate change.

## 2. Conceptual framework

### 2.1. Making hyperbolic equations efficient

To define an efficient mathematical equation for the hyperbola, we started with its Cartesian formulation, shown in equation 1. This corresponds to a hyperbola centred at the origin, where the x- and y-axes are the axes of symmetry. One branch of the hyperbola is above the x-axis, the other branch is below.

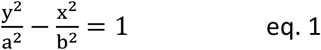

From this first equation, we can select one branch, expressed in equation 2.

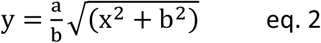

This gives one branch of the hyperbola that is symmetrical with the y-axis. To break this symmetry with the y-axis, a linear function can be added. Finally, to translate the hyperbola vertically, a constant can be added; to translate it horizontally, the x value can be changed to x minus a constant. This leads to the following equation (eq. 3), also called the “bent-hyperbola model” (Mesnil and Rochet, 2010; Watts and Bacon, 1974).

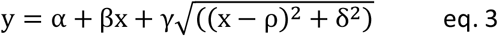

In this equation, the five parameters α, β, γ, ρ, and δ completely define the hyperbolic form. However, they are not easy to use, because they are not linked to graphically readable features. Cunningly chosen transformations of these five parameters can be applied to define five other parameters, the meaning of which can be read directly from the graph. These transformations produce a more complex mathematical equation, but one that is much more useful for interpreting the parameters. Furthermore, the new parameters are decorrelated from one another, which eases the convergence in the modelling process.

Equation 4 presents the result after transforming the parameters, without specifying all the steps required to achieve the process.

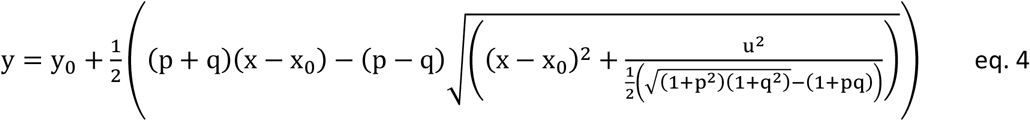

This equation is only valid for p ≠ q, otherwise the denominator inside the square root would equal zero. In all other cases, the denominator remains strictly positive, as demonstrated in the Supplementary Material section (S1, §A.1), making equation 4 similar to equation 3. Although complicated, this formulation is very useful because the parameters are meaningful (Figure 1): the parameters p and q are the slopes of the first and second asymptotes; x_0_ and y_0_ are the coordinates of the intersection between the two asymptotes; and u is the Euclidean distance between this intersection and the summit of the hyperbola. The equations for the asymptotes and bisectors are provided in the Supplementary Material section (S1, §A.2)

**Figure 1:**
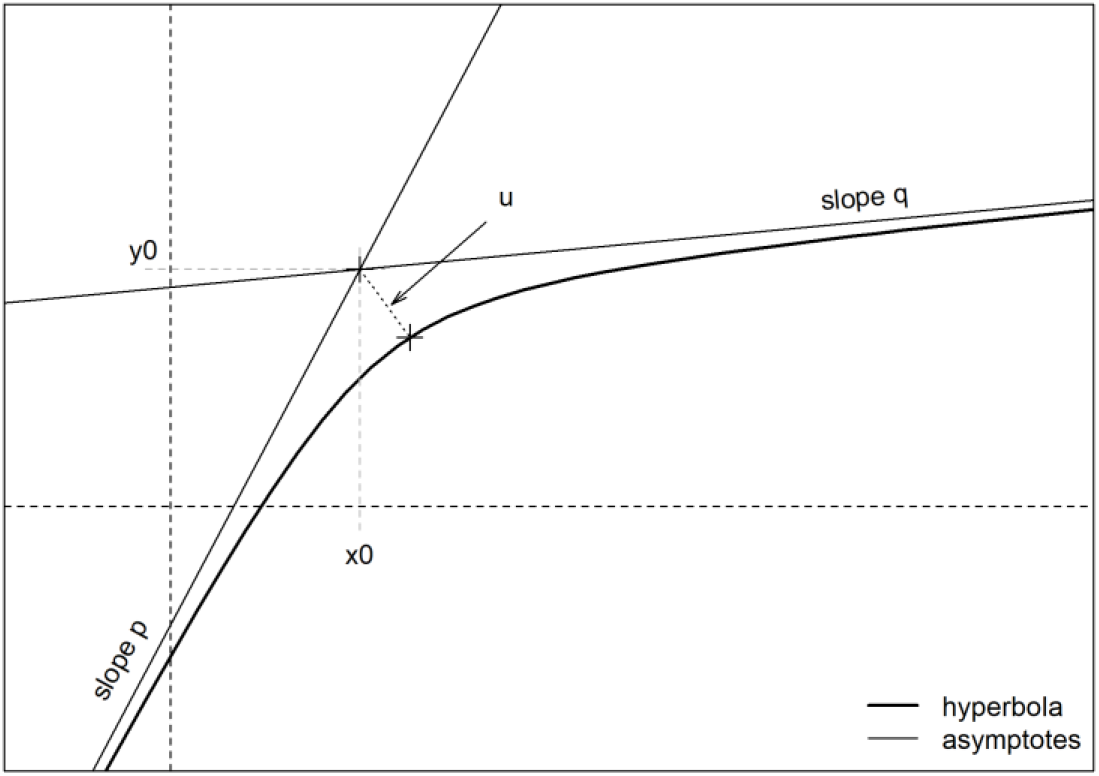
Meaning of the five parameters of the hyperbola defined with equation 4. The parameters p and q denote the slopes of the two asymptotes; the coordinates (x₀, y₀) specify the point where the asymptotes intersect; and u quantifies the distance between this intersection point and the summit of the hyperbola.

Figure 1 illustrates the fact that the five parameters are indeed decorrelated; for example, it is possible to change the slopes of the asymptotes independently, and without changing the position of the intersection (x_0_, y_0_). Figures B1 in Supplementary Material S1 provide several illustrations of changes in the parameters.

### 2.2. Using hyperbolic equations in model calibration

It is possible to calibrate models with the hyperbola described in equation 4 with non-linear fitting procedures such as the *nls* or *gnls* function in R. Supplementary Materials S2 and S3 provide the R code and the Python code that define the hyperbolic function, as well as an example plot of the function and an example of a non-linear model fitted with the hyperbolic function.

In equation 4 of the hyperbola, there is already an intercept (parameter y_0_). In order to add a variable in a model that already has an intercept, this parameter must be set to 0, otherwise the model would contain two distinct intercepts. In practice, this requirement means that only four extra parameters are needed to introduce a new variable with a hyperbolic form.

Furthermore, in most forest studies, the curvature of the hyperbola is small compared to the remaining variability in the response. Therefore, the distance u is generally small and non-significant. In this case, we recommend fixing u to a value different from zero but which remains small compared to the residual variance. Indeed, with u = 0, the hyperbola becomes equivalent to a two-segment form, and this creates a discontinuity in the derivative and makes it difficult for the algorithms to converge.

This reduces the hyperbola to a three-parameter function: one for the first slope, one for the second slope, and one for the x-position of the intersection of the asymptotes.

Finally, in some cases common in forestry, the effect of a variable on a process only begins (or saturates) at a given threshold value. In these cases, the corresponding slope can be set to zero, resulting in a two-parameter function.

## 3. Case study: hyperbolic response to climate in forest growth models

In this section, we used the growth modeling of 18 tree species in France with National Forest Inventory data as a case study. In particular, we used hyperbolic functions to fit the influence of climatic variables on growth where relevant.

### 3.1. Growth model framework

We based our new models on pre-existing model framework used for the Salem simulator (Aussenac et al., 2021; Vallet et al., 2021), which has proved its relevance through validation on long-term data (Mahnken et al., 2022). We modelled the basal area increment in forest stands over five years. Basal area increment depends on factors related to forest management, stand development stage and stand environmental conditions. The models are of the multiplicative type, as in equation 5 below,

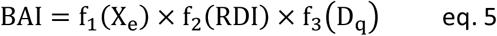

where BAI is the basal area increment of the stand over 5 years (m^2^/ha/5yr), f_1_ is a function of a set of environmental variables X_e_ (such as soil, topographic and climate variables), f_2_ is a function of stand density (RDI, see for example Aussenac et al., 2021; del Río et al., 2016) and f_3_ is a function of mean quadratic diameter D_q_, which is a proxy for stand development stage.

Based on results from former studies (Aussenac et al., 2021), we used identical forms for functions f_2_ and f_3_ for all 18 species. For these two functions in the model, only the parameters change among species. However, the X_e_ variables inside function f_1_ are not the same for all the tree species, because the trees species are not influenced by the same environmental factors. Since the effect of the X_e_ variables can be non-linear, we can use functions of these variables, including quadratic forms or hyperbolas (with fixing the u parameter to 0.1, because of the high variability of the data compared to the curvature). The general formulation of the model thus becomes

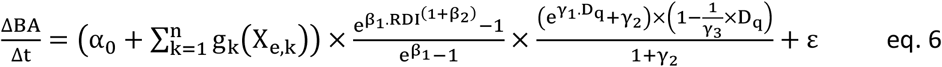

where α_0_, β_1_, β_2_, γ_1_, γ_2_, and γ_3_ are parameter to be fitted. To select the X_e_ variables and the g_k_ functions, we followed an ascending stepwise process, based on AIC diminution (Akaïke Information Criterion), fitting the complete model at each step. Once a variable was selected, variables with a similar ecological meaning could no longer be selected. To this end, we have categorised the climatic variables into three types: variables representing the length of the growing season (sum of growing degree days, minimum winter temperatures), summer constraints (summer water deficit, maximum summer temperatures, summer water balance), and water supply variables (annual precipitation, precipitation during the growing season). Only one variable from each category could be included in the model (see paragraph 3.2.1 for variable description). We also attempted to incorporate interactions among the variables X_e_ in the models, but this did not lead to any improvement. Finally, ε is the residual term, which follows a normal distribution. As the data showed heteroscedasticity, we added a *varPower* variance model, which increased with increasing fitted values (Pinheiro and Bates, 2006).

The growth model described in equation 6 has several interesting properties. Function f_2_ equals 1 when D_q_ equals 0, i.e., when the increment is known to be maximal. Function f_3_ equals 1 when RDI equals 1, i.e., when the stand is as its maximal cover capacity. As a consequence, the f_1_ function defines a site index corresponding to the maximum increase in basal area, obtained when diameter is minimal and density maximal.

We fitted one model for each of the 18 tree species using *gnls* function of the R package *nlme* according to the generalized least squares method (Pinheiro and Bates, 2006).

### 3.2. Data

#### 3.2.1. National Forest Inventory data

We fitted our growth models with data from the 2005 to 2022 surveys of the French National Forest Inventory (NFI) (IGN, 2024). The NFI is a systematic survey of the French forest, based on a 1km-by- 1km grid. Every year, one twentieth of the grid is selected, leading to more than 100,000 plots measured since 2005. A full description of the data is provided in Dalmasso et al. (2014). The field plots, where a sample of small, medium and large trees are measured, are made up of three concentric discs with radii of six, nine and 15 meters. The selection of trees across all size categories provides a comprehensive representation of the stand. The measurements used in our study were tree species, circumference and radial increments over the last 5 years, taken from tree cores (Dalmasso et al., 2014).

The NFI also measures a range of site features such as soil depth, stone content, soil texture, type of bedrock, etc. A floristic survey is carried out on the 15-metre-radius disc, making it possible to estimate other soil characteristics such as pH or the carbon to nitrogen ratio with bioindication models (Pinto et al., 2016).

Finally, thanks to the coordinates of the NFI plots and climate digital maps available at a 1-km resolution (Piedallu et al., 2013, data available at https://silvae.agroparistech.fr/), we estimated the 30-year mean monthly values for precipitation, minimum temperature, mean temperature, and maximum temperature for each site. In mountainous areas, a 1-km resolution can generate significant error due to the strong correlation between temperature and altitude. We corrected for this effect by calibrating temperature gradients with altitude (also called lapse rates), as in Combaud et al. (2024). Finally, we used precipitation and corrected temperatures to calculate climatic variables such as potential evapotranspiration (PET), water deficit (WD), climate water balance (CWB) and the annual sum of growing degree days (SGDD, defined as the sum of temperatures above 5.5 degrees Celsius over one year). In addition to annual means, we calculated the seasonal means of these climatic variables - spring (March to April), summer (June to August), fall (September to November) and winter (December to February), as well as values for the growing season (March to August). A description of all environmental variables used in the models is provided in Supplementary Material S1, section C.2.

#### 3.2.2. Plot selection

We selected pure even-aged forest plots from the NFI database. As the basal area increment was recorded from tree cores, we discarded plots that had been thinned in the last 5 years to avoid bias due to removed trees. Pure stands were those where the sampled trees were all of the same species.

Finally, we modelled the growth of all the tree species for which we had more than 50 plots. We ended up with a database of 8,330 plots representing 18 species (Table 1).

**Table 1:** Description of the calibration database

| Species | Abbr. | N plots | Mean diameter (cm) |  |  | Mean temperature (°C) |  |  | Precipitation (mm/year) |  |  |
| --- | --- | --- | --- | --- | --- | --- | --- | --- | --- | --- | --- |
|  |  |  | min | mean | max | min | mean | max | min | mean | max |
| *Abies alba* | Ab. al. | 326 | 5.7 | 29.7 | 84.1 | 4.8 | 8.9 | 12.2 | 715.9 | 1077.9 | 1917.2 |
| *Betula pendula* | Be. pe. | 65 | 5.2 | 16.8 | 48.9 | 6.7 | 10.7 | 13.1 | 674.2 | 921.7 | 1651.5 |
| *Carpinus betulus* | Ca. be. | 89 | 5.2 | 19.7 | 47.5 | 9.6 | 11.0 | 14.0 | 603.1 | 874.0 | 1262.1 |
| *Castanea sativa* | Ca. sa. | 195 | 5.1 | 22.7 | 98.5 | 9.0 | 12.0 | 15.6 | 638.8 | 1031.4 | 1760.7 |
| *Fagus sylvatica* | Fa. sy. | 797 | 5.1 | 31.7 | 117.0 | 5.3 | 9.5 | 14.3 | 675.0 | 1130.9 | 1974.3 |
| *Fraxinus excelsior* | Fr. ex. | 108 | 5.2 | 16.8 | 68.4 | 8.1 | 11.3 | 14.7 | 638.1 | 901.5 | 1442.0 |
| *Larix decidua* | La. de. | 166 | 5.5 | 28.5 | 66.7 | 1.6 | 5.6 | 11.2 | 733.1 | 913.7 | 1378.8 |
| *Picea abies* | Pi. ab. | 659 | 5.1 | 25.5 | 67.3 | 3.1 | 8.7 | 13.1 | 681.6 | 1177.4 | 1979.4 |
| *Pinus halepensis* | Pi. ha. | 251 | 5.6 | 22.7 | 56.2 | 11.5 | 14.4 | 16.4 | 548.2 | 699.5 | 1097.2 |
| *Pinus nigra laricio* | Pi. la | 305 | 5.0 | 22.0 | 77.7 | 7.2 | 11.5 | 14.5 | 630.6 | 936.7 | 1867.6 |
| *Pinus nigra nigra* | Pi. ni. | 219 | 5.1 | 19.8 | 47.5 | 5.9 | 10.6 | 15.0 | 662.5 | 942.5 | 1645.8 |
| *Pinus pinaster* | Pi. pi. | 1459 | 5.0 | 22.9 | 65.5 | 8.3 | 13.5 | 15.7 | 619.6 | 952.6 | 1603.5 |
| *Pinus sylvestris* | Pi. sy. | 985 | 5.1 | 22.5 | 63.9 | 4.4 | 9.6 | 15.1 | 651.4 | 941.1 | 1777.3 |
| *Pseudotsuga menziesii* | Ps. me. | 661 | 5.1 | 26.4 | 70.1 | 6.6 | 10.4 | 13.5 | 644.8 | 1012.6 | 1670.0 |
| *Quercus petraea* | Qu. pe. | 826 | 5.0 | 28.3 | 90.8 | 7.8 | 11.3 | 14.0 | 596.4 | 814.2 | 2039.1 |
| *Quercus pubescens* | Qu. pu. | 451 | 5.6 | 19.5 | 64.5 | 7.7 | 12.4 | 16.0 | 637.6 | 908.2 | 1668.6 |
| *Quercus robur* | Qu. ro. | 713 | 5.1 | 30.6 | 108.4 | 7.7 | 12.0 | 15.2 | 619.9 | 910.4 | 1875.6 |
| *Robinia pseudoacacia* | Ro. ps. | 55 | 5.5 | 13.8 | 58.5 | 9.6 | 12.3 | 14.4 | 678.0 | 908.3 | 1436.4 |

### 3.3. Results

We fitted our growth models in even-aged pure stands for 18 tree species. The full description of the parameters and the figures of the models’ residuals against the fitted values are provided in Supplementary Material S1, section C.

#### 3.3.1. Effects of climate on growth dynamics

The stepwise selection highlighted the importance of three types of climate variable on basal area increment. The Sum of Growing Degree Days and the winter minimum temperature reflect the start of the vegetation period, and by extension, its duration. Summer water deficit, summer maximal temperature and summer climatic water balance represent the summer constraints to growth. Annual precipitation reflects the global water supply. Figure 2 shows the effect of these variables, with the left-hand column of graphs representing the effect of the growing season and precipitation, and the right-hand column of graphs representing summer constraints. In these figures, the partial effects are shown for one climatic variable, all other environmental variables being set to their mean value, or to their modal value for categorical or Boolean variables.

**Figure 2:**
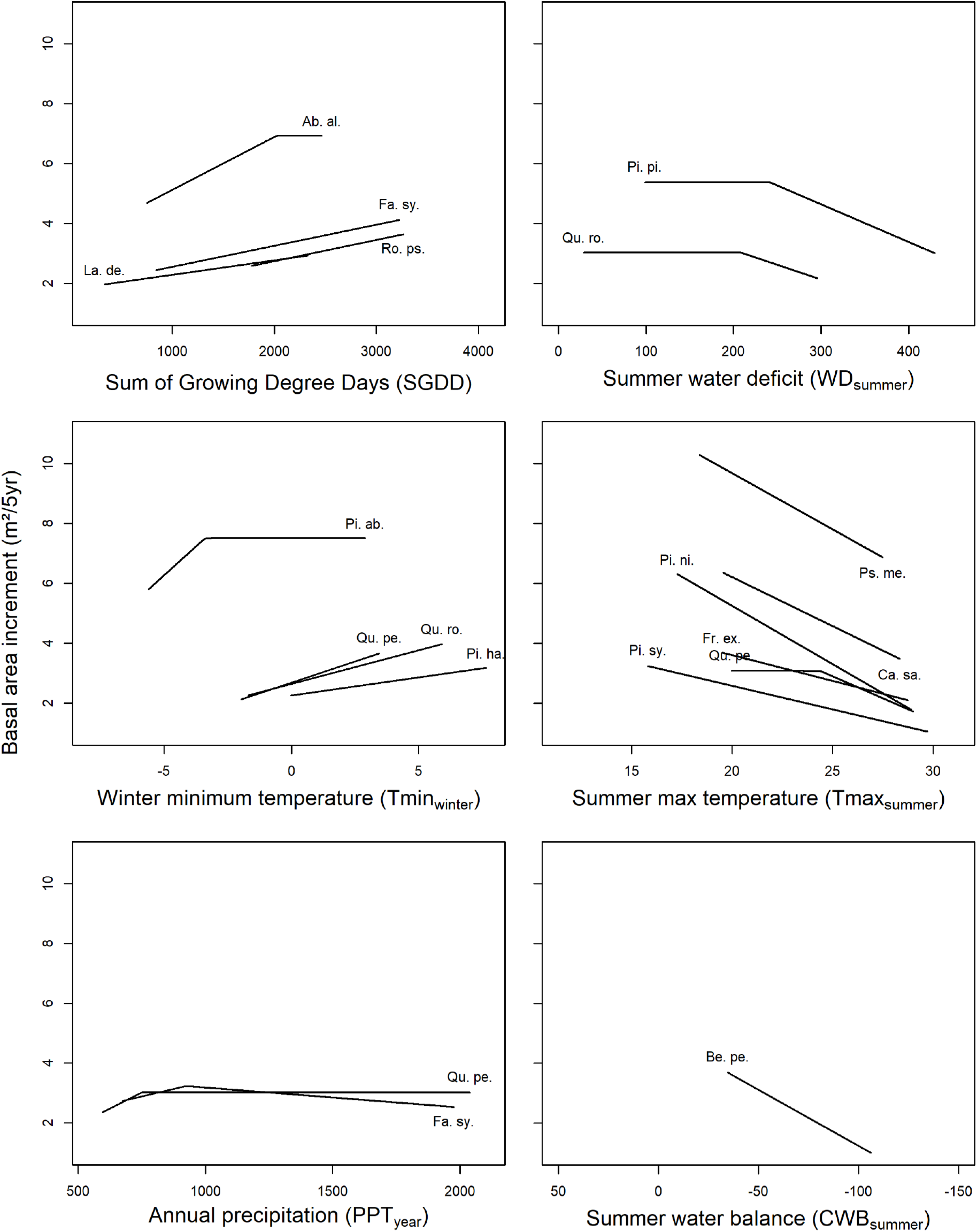
Curves representing all partial climate effects for the different species, holding other variables constant. All other environmental variables were set to their mean values for each species, or to their modal value for categorical or Boolean variables. The mean quadratic diameter was set to 25 cm, and the RDI to 0.7. The range corresponds to that of the species dataset. The CWB graph was x-axis reversed to show constraints, as for the other summer graphs. Hyperbolic responses exhibit a shallow curvature, which reflects the low value assigned to the u parameter. The abbreviations of the names are indicated in table 1.

We used the hyperbolic rather than linear form of climatic variables in seven cases out of 19. For the SGDD for Abies alba, the winter minimum temperature for Picea abies and annual precipitation for Quercus petraea and Fagus sylvatica, the hyperbola made it possible to detect a positive effect at low values, followed by a constant or slightly negative effect for higher values. In the case of Quercus robur and Pinus pinaster, the negative effect of water deficit started only at strong levels of constraint (at relatively, 207 mm and 241 mm for the three months). For Quercus petraea, the summer maximum temperature negatively impacted growth only above 24.4°C, whereas the negative effect was linear for all other species (when significant) over the entire range of maximum summer temperatures.

#### 3.3.2. Effects of other environmental conditions

Thanks to the wealth of information provided by NFI data, we were able to evaluate the influence of various topographical and edaphic factors on stand growth. The influence of these variables aligns with our understanding of forest species ecology (Table 2). The carbon-to-nitrogen ratio negatively affected growth in 12 out of the 18 models, demonstrating the importance of soil nitrogen richness for the selected species. Soil water holding capacity (SWHC) had a positive effect on growth for nine species. pH was also an important factor, involved in eight models. pH took the form of a quadratic equation (for four species), either reaching a maximum at values from 4.5 to 5.5, or having only a negative effect. We observed a negative influence of limestone bedrock for five species and of rocky outcrops for four species. Finally, residual geographical effects (Greco) were observed for six species.

**Table 2:**
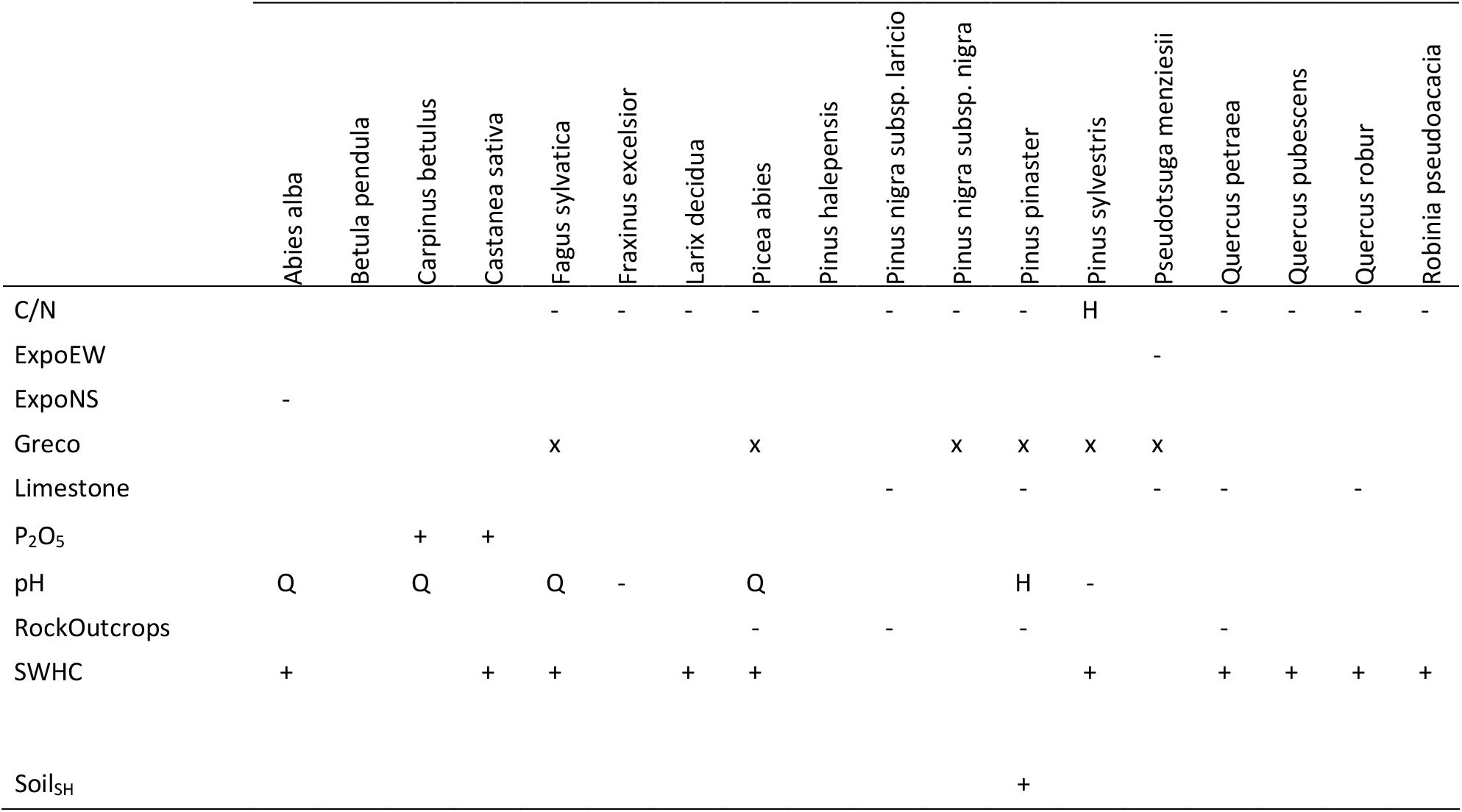
Summary of non-climatic environmental factors. The “+” or “-” symbols indicate the sign of the effect. The letter “H” indicates a hyperbolic effect, “Q” a quadratic effect and “x” an effect that can be positive or negative depending on the modality. A detailed description of the models is provided in Supplementary Material S1, section C.

#### 3.3.3. Effects of dendrometrical conditions

Figure 3 illustrates the influence of stand dendrometric variables on species growth. Figure 3a shows that growth increases with stand density (RDI), reaching maximum growth when RDI equals 1. Broadleaves species have a differentiated density effect, while all pines exhibit a similar effect, as do the other coniferous species. Figure 3b shows that growth is maximal for small stand diameters, then decreases to zero. The diameter value at which the curve intersects the x-axis corresponds to parameter γ_3_ of equation 6, and represents the asymptotic value of D_q_ of the species as time increases. γ_3_is therefore the maximum mean quadratic diameter value a species can reach.

**Figure 3:**
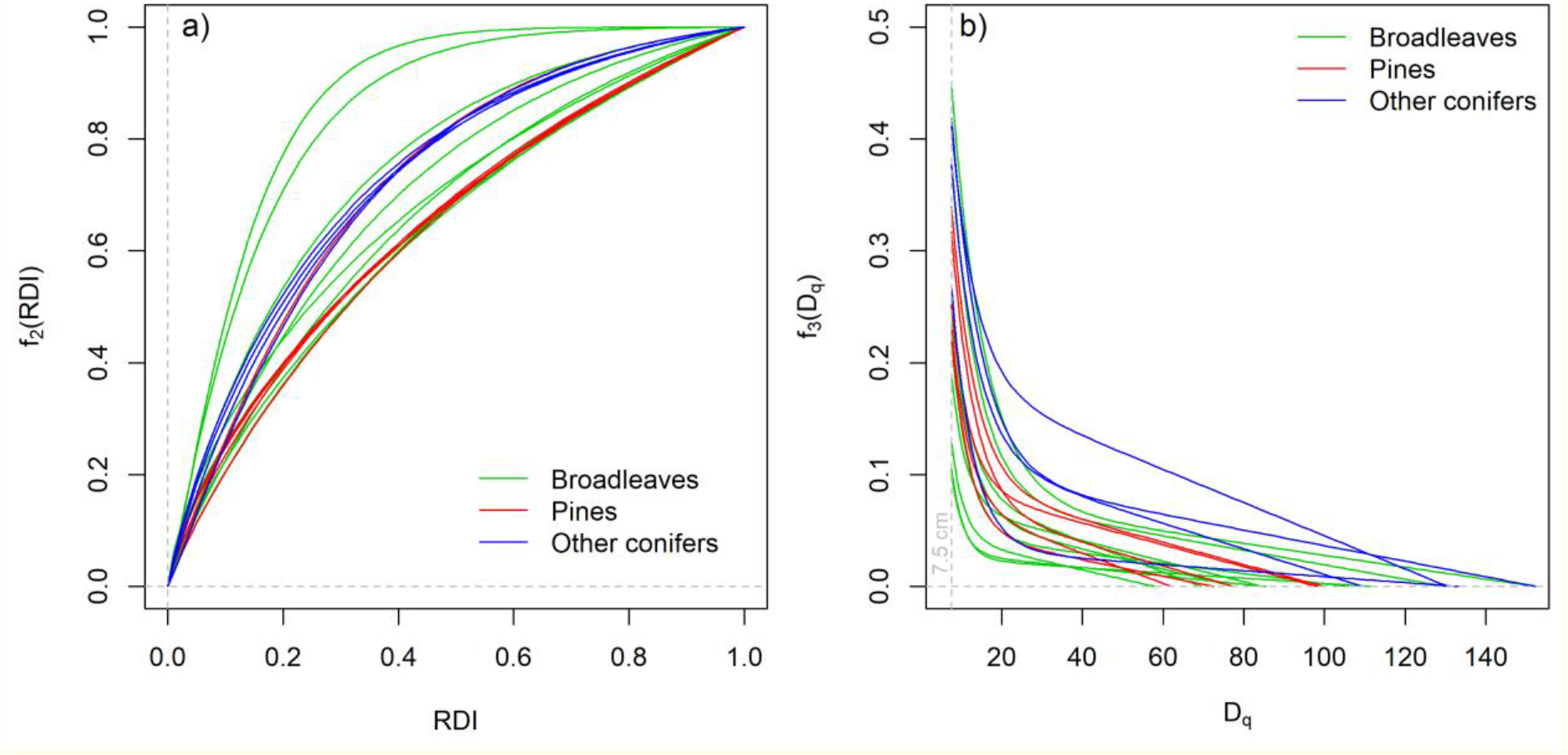
Partial effects of stand dendrometric variables on stand growth for the final fitted models as described by equation 6: a) effect of density index (RDI), and b) effect of mean quadratic diameter (D_q_). Each line represents one of the 18 fitted species. Functions f_2_ and f_3_ and reducers, and standardized between 0 and 1.

## 4. Discussion

### 4.1. Interests and limits of the hyperbolic formulation

In this article, we present a five-parameter mathematical formulation of the hyperbola. The five parameters are self-explanatory, as they correspond to easily understandable characteristics such as the slope of the two asymptotic branches, or the x and y position. In practice, when modelling processes in ecology, often three parameters suffice, or even only two when the effect of a variable is non-significant (i.e., one slope equals zero).

The main advantage of the hyperbolic form is that it makes it easy to take into account the effects of a variable that is close to segmented effect. For example, in the model for the sessile oak, we identified an effect of annual precipitation, but only for values below approximately 800 mm per year. Above this threshold, there is no longer any effect. Figure 4 illustrates the usefulness of the hyperbola in this case. The plot on the left shows the normalized residuals as a function of precipitation for a model similar than the final model, but before the precipitation variable was included. The plot on the right shows the model residuals as a function of precipitation once this variable has been added. The hyperbolic form made it possible to account for the segmented response to precipitation and to obtain a valid residual structure in the final model.

**Figure 4:**
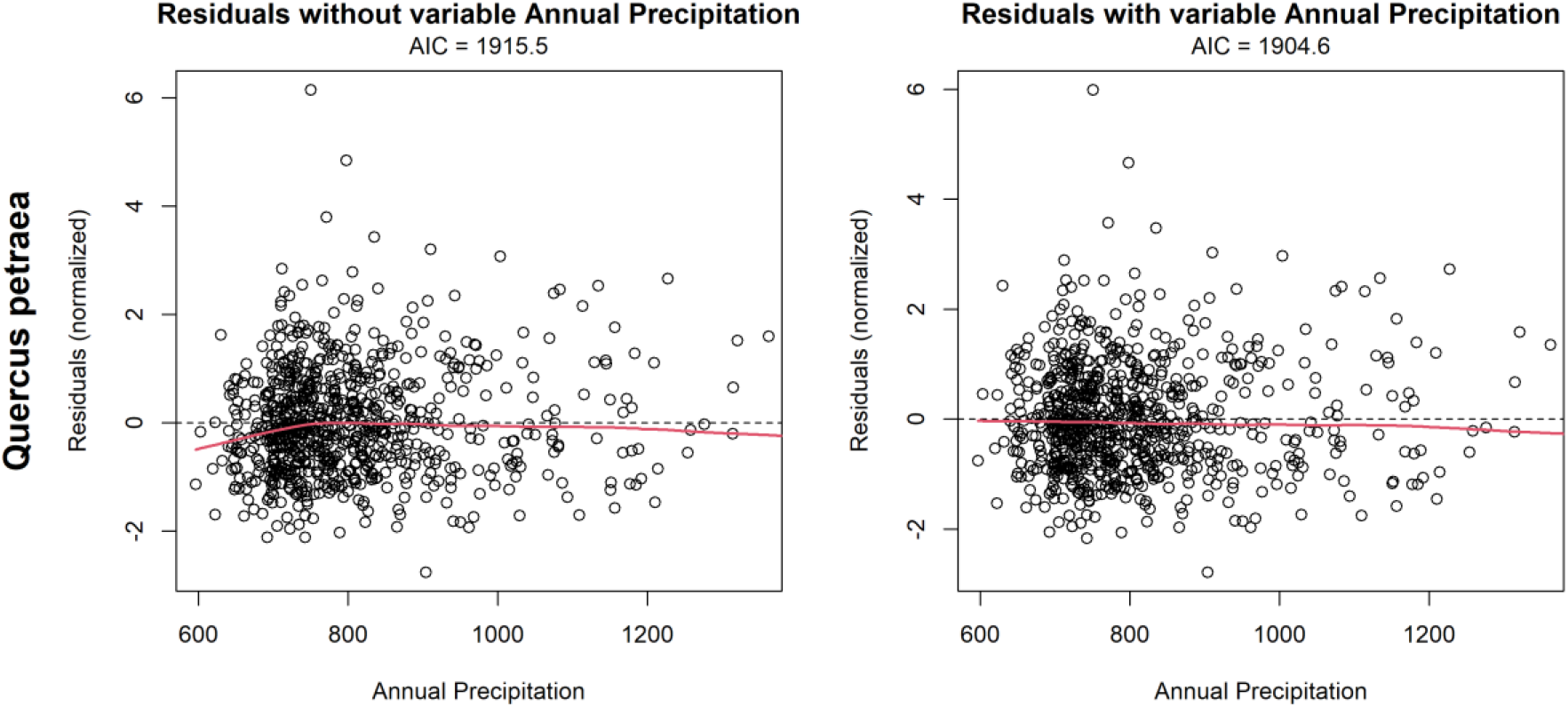
Normalized residuals against Annual Precipitation for the Quercus petraea model. On the left side, the residuals are for a model before the inclusion of the Annual Precipitation variable. On the right side, the residuals are for the model with this inclusion of the Annual Precipitation variable, and also corresponds to the final model. The red curves are *lowess* smoothing curves.

We can observe on this graph a substantial residual variability relative to the magnitude of the precipitation effect. This outcome was expected, as the influence of environmental variables on growth is of second order compared with dendrometric variables—especially RDI—and because we rely on observational NFI data rather than controlled experimental data. Nonetheless, incorporating precipitation improves the model by roughly 11 AIC points, demonstrating that even modest effects can be calibrated using this hyperbolic equation.

The corresponding figures are provided in the Supplementary Materials S1, §D, for the seven other model cases in which we observed hyperbolic effects of the climatic variables.

Alternative methods such as segmented curves exist to account for this type of response to environmental factors. However, compared to these forms, the hyperbola has the major advantage of avoiding discontinuity in the first derivative, which eases convergence when fitting the models.

Two attentions should be taken when hyperbolic functions are used. The first is that, in many modelled systems, small effects can be identified at the extremes of the range of values covered by the explanatory variables. It may be tempting to use a hyperbolic function to account for these effects, even though they may be mere artifacts at the edge of the range covered by the data, and only linked to response variability. Furthermore, since it is easy to add a variable with a hyperbolic form, it may be tempting to add this type of response to several variables simultaneously. This can result in over-parameterized models – even when metrics penalizing the addition of variables such as AIC – or models with excessive correlation between variables, but in a less obvious way than in a more traditional linear adjustment. We therefore recommend parsimony when adding parameters and checking the consistency of the modelled results with actual knowledge (Box, 1979).

The second attention concerns fitting parameter u, representing the curvature of the hyperbola, from equation 4. This curvature can be small compared to the variability of the response. In such cases, it may be too complicated to fit u so that the model converges. This issue can be solved by fixing u at a small value compared to the standard deviation of the data. If the value is low compared to the standard deviation, the choice of the value itself is irrelevant as it will only have a very marginal impact on the other parameters. However, the value must always be different from zero to avoid discontinuity in the derivative, which would prevent the model from converging.

### 4.2. Forest growth models

#### 4.2.1. Climate effects on growth

As expected, the variables SGDD and Tmin_winter_ had a positive effect on growth. The effect was significant for eight species out of 18, thus corroborating previous results (Albert and Schmidt, 2010; Antón-Fernández et al., 2016; Seynave et al., 2008). The climate variables used in our study were thirty-year averages for the period 1991-2020. Their variations are linked to the location of the NFI plots, and not to temporal variations in the climate. SGDD and Tmin_winter_ enabled us to characterize more or less warm sites, and therefore to represent the extent of the growing season. The effect of SGDD and Tmin_winter_ also corresponds to multiple findings in the literature linking elevation and a decrease in productivity (see for example Brandl et al., 2018; Aussenac et al., 2021 in temperate forests; or Muller-Landau et al., 2021 in tropical forests), interpreted in these articles by a decrease in temperature for higher elevations. In our study, we deliberately excluded elevation from the possible explanatory variables, and preferred to use direct climate variables such as mean temperatures (or SGDD), summer temperatures or precipitation. In order to run projections in a changing climate, it is more appropriate to use direct climate variables rather than proxies such as elevation, which do not change over time. The hyperbolic functions enabled us to disentangle the effects of these multiple variables.

The effect of water availability was different depending on the species. For Quercus petraea and Fagus sylvatica, we found a positive effect of annual precipitation on basal area increment, which corroborates previous studies (Michelot et al., 2012). However, this effect occurred only at low values of precipitation (below 750 mm of rainfall for Quercus petraea and below 920 mm for Fagus sylvatica), becoming null or slightly negative above these values. When precipitation is too low growth will decrease, but above a given level, water is no longer limiting and variations in precipitation no longer have an effect. For Quercus robur, Pinus pinaster, and Betula pendula, the water availability effect was linked to summer climatic constraints such as summer water deficit or summer water balance. Once again, the effect of summer water deficit on Quercus robur and Pinus pinaster became null when the water deficit for the three summer months fell below approximately 200mm, and negative for higher values. For Betula pendula, the decrease in growth with increasing summer constraint was due to the effect of the summer water balance. The effect of water availability on species growth is well documented in the scientific literature on forestry, showing both a positive effect of the overall annual water balance and a negative effect of summer drought (see, for example, Camarero et al. (2025) for a recent article dealing with Quercus robur and Pinus pinaster).

Maximum summer temperature emerged as the predominant summer constraint, affecting the greatest number of species (6 out of 18), thereby exerting a stronger influence than drought stress. Maximum temperature has already been identified as an important driver of species climate-growth patterns (Serrano-Notivoli et al., 2025) and ring width (Arsalani et al., 2015). This relationship arises from the role high temperatures play in controlling the cessation of cell enlargement (Guada et al., 2020).

Overall, we identified an effect of summer climatic constraints for nine species out of 18. For these species, climate change is becoming a serious impediment (Cheaib et al., 2012), even though they may have experienced an increased growth rate during the 20^th^ century due to a reduction in thermal constraints (Combaud et al., 2024; Socha et al., 2021).

#### 4.2.2. Environmental and dendrometrical effects on growth

The effects of the other environmental variables we considered correspond to well-known effects on forest growth (Piedallu et al., 2016). One of the main factors is the carbon to nitrogen ratio (Table 2). We selected this variable for twelve species, always with a negative effect. A decrease in the carbon to nitrogen ratio corresponds to an increase in soil nitrogen availability, and therefore to higher soil fertility (Albert and Schmidt, 2010; Sharma et al., 2012). Soil pH was identified as another important factor, included in the model for seven species. For four species, the pH effect followed a concave quadratic form. The way we defined the quadratic form (i.e., a × (pH − b)^2^) allowed us to easily identify the optimum pH for a given species. This was simpler than a hyperbolic form, thus showing the benefits of defining the appropriate mathematical form of a model. Here, the b parameter ranged from 4.35 to 5.55 depending on the species, in accordance with the known optima. For instance, Piedallu et al. (2016) report that Carpinus betulus occurs across a soil-pH interval of 3.3–6.8; in our analyses the species showed an optimal pH of 5.55. Likewise, in Piedallu et al. (2016) Picea abies has a pH range of 3–7.1, with an optimum of 4.35 in our study. Soil water holding capacity was selected in ten of our models. This was expected since the variable reflects the maximum amount of water in the soil available to the stand (Roces-Diaz et al., 2025). Other variables such as rock outcrops or limestone bedrock were also identified as constraints for some species, and correspond to previously published results for the same species (Aussenac et al., 2021). Finally, geographical effect remained, but we could not identify the underlying biogeophysical variables. However, the effects remained small.

For dendrometrical variables, the stand density effect, expressed here by the RDI, correspond to results from the literature (Allen and Burkhart, 2019; Trouvé et al., 2019), although the precise form of the relationship is still under debate. As expected, stand productivity increased with stand density in our models. Finally, the other dendrometrical factor taken into account was the effect of the mean quadratic diameter, as identified in previous studies (Toïgo et al., 2015; Vallet and Perot, 2018; Vallet and Pérot, 2011). Compared to our previous studies, we improved the model by adding a parameter (γ_3_) corresponding to the maximum diameter a species can reach, which increased model flexibility. For the 18 species, this parameter ranged from 57.9 cm for Betula pendula to 152.5 cm for Abies alba and Fagus sylvatica (cf. Supplementary Materials S1, §C3a., C3b and C3e). This is consistent with the maximum diameters that can be observed in the NFI data for these species.

## 5. Conclusion

In this study, we developed a mathematical formulation of the hyperbola that enables a clear decoupling of its defining parameters while providing practical interpretability. The parameters correspond to the slopes of the two asymptotes, the position in the plane, and the radius of curvature. This formulation facilitates modelling ecological processes that exhibit responses having such a hyperbolic behaviour.

We illustrated the possibility of this new formulation by applying our hyperbolic equation to forest growth modelling. The hyperbolic equation made it possible to capture climatic effects occurring above or below specific thresholds — for instance, maximum summer temperature or summer soil water deficit — for seven cases out of or 18 forest species. In the context of climate change, it is particularly important to properly account for non-linear responses to environmental drivers and, in particular, to identify summer climate tipping points.

We provide ready-to-use R and Python codes for our hyperbolic equation in Supplementary Materials S2 and S3.

## Supporting information

R Script for hyperbola

Python Script for hyperbola

## Acknowledgements

This research was conducted as part of the MELBAC project, which received financial support from France-Bois-Forêt, the national interprofessional organization for the forest and timber industry (grant number 24RD1986). The author also thanks the French National Forest Inventory for freely providing the data.

## Data, statistical scripts, and code availability

All the data used in this article and all the scripts to calibrate the models and generate the figures and tables (including for the supplementary material) are available in the following archive: https://doi.org/10.5281/zenodo.18805407

