## Supplementary Material S1 for "A hyperbolic equation for modelling non-linear responses of forest tree growth to environmental factors"

### A. Mathematical considerations

#### 1. Demonstration of the positivity of the numerator in equation 4

In equation 4, to meet the generic formulation of the hyperbola (equation 3), the term in the numerator must be positive.

This numerator is given by the following equation (see equation 4 of the main manuscript):

$$\frac{1}{2}\left( \sqrt{\left( 1+p^{2} \right)\left( 1+q^{2} \right)}-\left( 1+pq \right) \right)$$

We can calculate the difference of the square value of the two main terms of this equation:

$${\sqrt{\left( 1+p^{2} \right)\left( 1+q^{2} \right)}}^{2}-\left( 1+pq \right)^{2}$$

$$=\left( 1+p^{2}+q^{2}+p^{2}q^{2} \right)-\left( 1+2pq+p^{2}q^{2} \right)$$

$$=p^{2}+q^{2}-2pq$$

$$=\left( p-q \right)^{2}$$

As this difference is a square value, it is positive. Therefore

$${\sqrt{\left( 1+p^{2} \right)\left( 1+q^{2} \right)}}^{2}\geq\left( 1+pq \right)^{2}$$

As the square root is a monotonically increasing function, we have

$$\sqrt{\left( 1+p^{2} \right)\left( 1+q^{2} \right)}\geq\left( 1+pq \right)$$

And then

$$\sqrt{\left( 1+p^{2} \right)\left( 1+q^{2} \right)}-\left( 1+pq \right)\geq0$$

Which demonstrates that the numerator is positive.

#### 2. Equation of the asymptotes and bisectors

For the hyperbola defined as in equation 4, the asymptotes and bisectors equations are as follow:

First asymptote:

$y=y_{0}+p\times\left( x-x_{0} \right)$

Second asymptote:

$y=y_{0}+q\times\left( x-x_{0} \right)$

First bisector:

$y=y_{0}+\frac{p.\sqrt{1+q^{2}}-q.\sqrt{1+p^{2}}}{\sqrt{1+q^{2}}-\sqrt{1+p^{2}}}\times\left( x-x_{0} \right)$

Second bisector:

$y=y_{0}+\frac{p.\sqrt{1+q^{2}}+q.\sqrt{1+p^{2}}}{\sqrt{1+q^{2}}+\sqrt{1+p^{2}}}\times\left( x-x_{0} \right)$

### B. Parameters’ variation of the hyperbola

The figure B1 shows the variation of form of the hyperbola as described in equation 4 when the five parameters vary. This illustrates the fact that the five parameters are decorrelated.

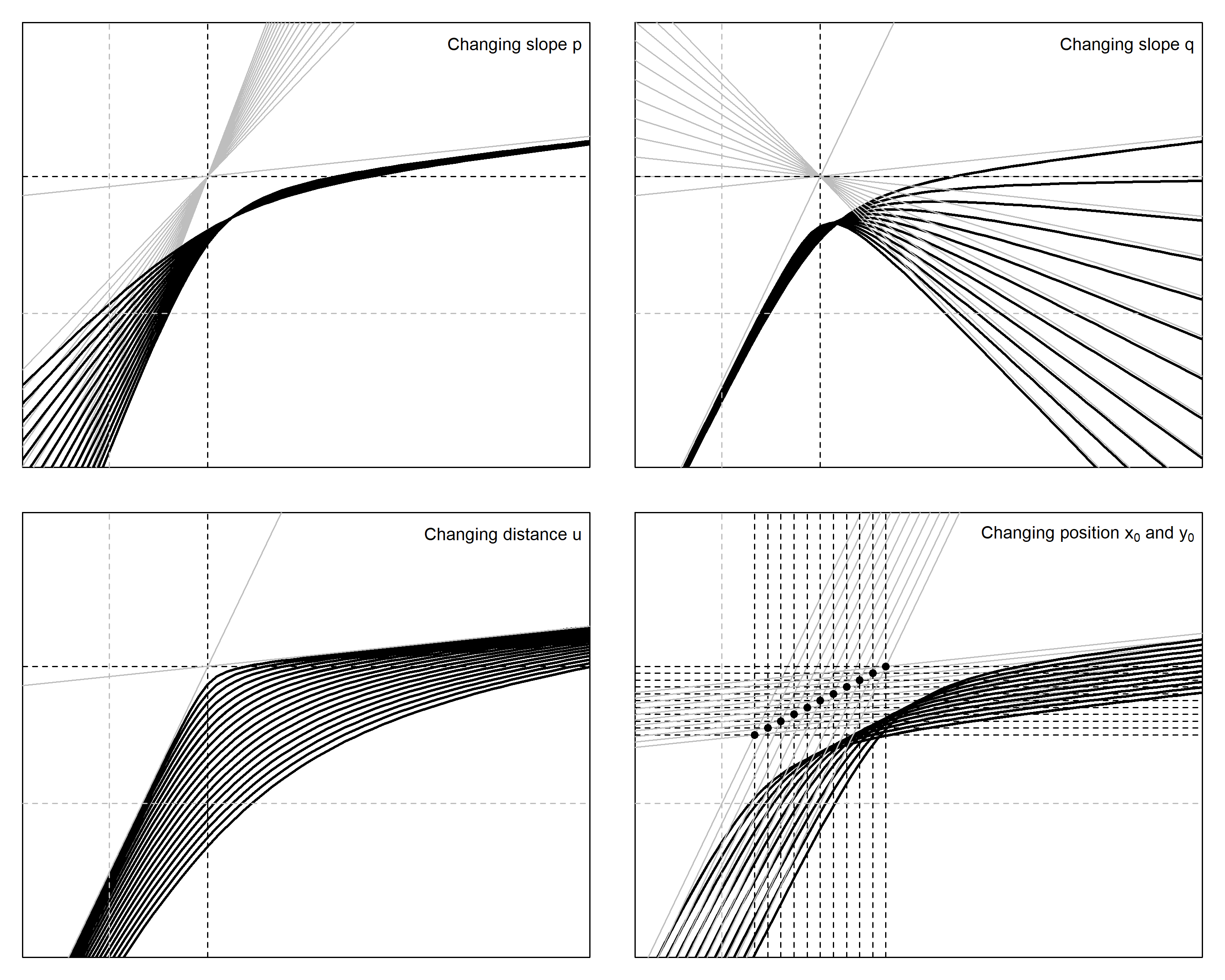

Figure B1. Variation of the hyperbola when one of the parameters vary.
From left to right, and top to bottom:

- variation of the first slope (parameter p)
- variation of the second slope (parameter q)
- distance between asymptotes intersection and the summit of the hyperbola (parameter u)
- position of the intersection (parameters x_0_ and y_0_)

### C. Description of the models for the 18 calibrated species

#### 1. Basal area increment model’s complete formula

The general form of the model is given by equation 5 of the main manuscript:

$$\frac{\Delta BA}{\Delta t}=f_{1}\left( X_{e} \right)\times f_{2}\left( \mathrm{RDI} \right)\times f_{3}\left( D_{q} \right)$$

The developed form is given by equation 6, which is reminded here:

$$\frac{\Delta BA}{\Delta t}=\left( \alpha_{0}+\sum_{k=1}^{n} g_{k}\left( X_{e,k} \right) \right)\times\frac{e^{\beta_{1}.\mathrm{RDI}^{\left( 1+\beta_{2} \right)}}-1}{e^{\beta_{1}}-1}\times\frac{\left( e^{\gamma_{1}.D_{q}}+\gamma_{2} \right)\times\left( 1-\frac{1}{\gamma_{3}}\times D_{q} \right)}{1+\gamma_{2}}+\varepsilon$$

As explained in the main text, the variables X_e_ and the function g_k_ differ between the species and were selected with a stepwise method. The other parts of the model are similar between species. For the presentation of the results below, only the function f_1_ corresponding to the potential is developed.

In all the species equations below, if a hyperbola is used, as the model already contains an intercept ($\alpha_{0}$), the parameter y_0_ is set to 0. Moreover, the distance “u” was fixed to 0.1. We simplified the notation as:

$$\mathrm{hyp}\left( X_{e}, p, q, x_{0} \right)=hyperbola\left( X_{e}, p, q, u=0.1, x_{0}, y_{0}=0 \right)$$

where hyp is the hyperbola function as defined in equation 4 (and provided in supplementary material S2 “Hyperbola.R”), X_e_ is the environmental variable, p is the first slope, q the second slope, and x_0_ the value of X_e_ for the crossing of the asymptotes.

#### 2. Involved environmental variables

The table below describes all the variables involved in the growth models.

| **Variable** | **Meaning** | **Unit** |
| --- | --- | --- |
| C/N | Carbon over Nitrogen ratio^[1]^ | -- |
| CWB | Climatic Water Balance | mm |
| ExpoEW | Sinus of plot aspect | -1 (West) to 1 (East) |
| ExpoNS | Cosinus of plot aspect | -1 (South) to 1 (North) |
| Greco | Geographical area (11 areas from A to K in France)^[2]^ | Boolean |
| Limestone | Limestone Bedrock | Boolean |
| P_2_O_5_ | Phosphorus content^1^ (log value from Duchaufour method) | -- |
| pH | Acidity^1^ | -- |
| PPT | Precipitation | mm |
| RockOutcrops | Rock Outcrops | % /10 |
| SGDD | Sum of Growing Degree-Days | °C |
| SWHC | Soil Water Holding Capacity | mm |
| Soil_SH_ | slightly hydromorphic soils | Boolean |
| Tmin | Mean of minimum temperature | °C |
| Tmean | Mean of mean temperature | °C |
| Tmax | Mean of maximum temperature | °C |
| WD | Water deficit | mm |

Some climatic variables are calculated over periods of time. These periods are provided in the following table:

| **Period** | **Months accounted** |
| --- | --- |
| Spring | March to May |
| Summer | June to August |
| Fall | September to November |
| Winter | December to February |
| Growing season | March to August |
| Year | Full year |

Therefore, for example, the notation Tmax_summer_ corresponds to the mean of maximum temperatures for months June to August.

#### 3. Models’ parameters

##### a. Abies alba

Function f_1_ for Abies alba is given by:

$$f_{1}\left( X_{e} \right)=\alpha_{0}+\alpha_{1}.\left( pH-\alpha_{2} \right)^{2}+\alpha_{3}.SWHC+\alpha_{4}.ExpoNS+hyp\left( SGDD, p=\alpha_{5}, q=0, x_{0}=\alpha_{6} \right)$$

**Table of parameters for Abies alba:**

| **Function** | **Corresponding factor** | **Parameter** | **Estimate** | **Std. Error** |
| --- | --- | --- | --- | --- |
| f_1_ | Intercept | $\alpha_{0}$ | 57.93 | 17.91 |
|  | pH | $\alpha_{1}$ | -5.49 | 1.88 |
|  | pH | $\alpha_{2}$ | 5.44 | 0.12 |
|  | SWHC | $\alpha_{3}$ | 0.16 | 0.06 |
|  | ExpoNS | $\alpha_{4}$ | -5.22 | 2.40 |
|  | hyp(SGDD): slope 1 | $\alpha_{5}$ | 0.0163 | 0.0072 |
|  | hyp(SGDD): x position | $\alpha_{6}$ | 2023.43 | 305.99 |
| f_2_ | RDI | $\beta_{1}$ | -2.730 | 0.776 |
|  | RDI | $\beta_{2}$ | -0.126 | 0.080 |
| f_3_ | D_q_ | $\gamma_{1}$ | -0.131 | 0.024 |
|  | D_q_ | $\gamma_{2}$ | 0.119 | 0.032 |
|  | D_q_ | $\gamma_{3}$ | 152.5 | 53.9 |

##### b. Betula pendula

Function f_1_ for Betula pendula is given by:

$$f_{1}\left( X_{e} \right)=\alpha_{0}+\alpha_{1}.\mathrm{CWB}_{\mathrm{summer}}$$

**Table of parameters for Betula pendula:**

| **Function** | **Corresponding factor** | **Parameter** | **Estimate** | **Std. Error** |
| --- | --- | --- | --- | --- |
| f_1_ | Intercept | $\alpha_{0}$ | 193.51 | 109.46 |
|  | CWB_summer_ | $\alpha_{1}$ | 1.45 | 0.86 |
| f_2_ | RDI | $\beta_{1}$ | -3.117 | 1.542 |
|  | RDI | $\beta_{2}$ | -0.084 | 0.089 |
| f_3_ | D_q_ | $\gamma_{1}$ | -0.301 | 0.064 |
|  | D_q_ | $\gamma_{2}$ | 0.050 | 0.026 |
|  | D_q_ | $\gamma_{3}$ | 57.9 | 5.7 |

##### c. Carpinus betulus

Function f_1_ for Carpinus betulus is given by:

$$f_{1}\left( X_{e} \right)=\alpha_{0}+\alpha_{1}.\left( pH-\alpha_{2} \right)^{2}+\alpha_{3}.P_{2}O_{5}$$

**Table of parameters for Carpinus betulus:**

| **Function** | **Corresponding factor** | **Parameter** | **Estimate** | **Std. Error** |
| --- | --- | --- | --- | --- |
| f_1_ | Intercept | $\alpha_{0}$ | 69.50 | 38.53 |
|  | pH | $\alpha_{1}$ | -8.80 | 6.80 |
|  | pH | $\alpha_{2}$ | 5.55 | 0.24 |
|  | P_2_O_5_ | $\alpha_{3}$ | 19.83 | 14.16 |
| f_2_ | RDI | $\beta_{1}$ | -2.464 | 1.284 |
|  | RDI | $\beta_{2}$ | -0.039 | 0.093 |
| f_3_ | D_q_ | $\gamma_{1}$ | -0.205 | 0.065 |
|  | D_q_ | $\gamma_{2}$ | 0.105 | 0.052 |
|  | D_q_ | $\gamma_{3}$ | 69.4 | 13.9 |

##### d. Castanea sativa

Function f_1_ for Castanea sativa is given by:

$$f_{1}\left( X_{e} \right)=\alpha_{0}+\alpha_{1}.SWHC+\alpha_{2}.P_{2}O_{5}+\alpha_{3}.\mathrm{Tmax}_{\mathrm{summer}}$$

**Table of parameters for Castanea sativa:**

| **Function** | **Corresponding factor** | **Parameter** | **Estimate** | **Std. Error** |
| --- | --- | --- | --- | --- |
| f_1_ | Intercept | $\alpha_{0}$ | 800.16 | 290.72 |
|  | SWHC | $\alpha_{1}$ | 0.81 | 0.38 |
|  | P_2_O_5_ | $\alpha_{2}$ | 116.84 | 53.55 |
|  | Tmax_summer_ | $\alpha_{3}$ | -18.87 | 8.52 |
| f_2_ | RDI | $\beta_{1}$ | -1.364 | 0.744 |
|  | RDI | $\beta_{2}$ | -0.086 | 0.097 |
| f_3_ | D_q_ | $\gamma_{1}$ | -0.324 | 0.037 |
|  | D_q_ | $\gamma_{2}$ | 0.027 | 0.008 |
|  | D_q_ | $\gamma_{3}$ | 111.8 | 9.1 |

##### e. Fagus sylvatica

Function f_1_ for Fagus sylvatica is given by:

$$f_{1}\left( X_{e} \right)=\alpha_{0}+\alpha_{1}.\left( pH-\alpha_{2} \right)^{2}+\alpha_{3}.C/N+\alpha_{4}.SWHC+\alpha_{5}.SGDD+hyp\left( \mathrm{PPT}_{\mathrm{year}}, p=\alpha_{6}, q=\alpha_{7}, x_{0}=\alpha_{8} \right)+\alpha_{9}.\mathrm{Greco}_{B}+\alpha_{10}.\mathrm{Greco}_{C}+\alpha_{11}.\mathrm{Greco}_{D}+\alpha_{12}.\mathrm{Greco}_{J}$$

**Table of parameters for Fagus sylvatica:**

| **Function** | **Corresponding factor** | **Parameter** | **Estimate** | **Std. Error** |
| --- | --- | --- | --- | --- |
| f_1_ | Intercept | $\alpha_{0}$ | 21.39 | 4.16 |
|  | pH | $\alpha_{1}$ | -0.87 | 0.33 |
|  | pH | $\alpha_{2}$ | 5.39 | 0.26 |
|  | C/N | $\alpha_{3}$ | -0.3421 | 0.0981 |
|  | SWHC | $\alpha_{4}$ | 0.1108 | 0.0224 |
|  | SGDD | $\alpha_{5}$ | 0.0073 | 0.0017 |
|  | hyp(PPT_year_): slope 1 | $\alpha_{6}$ | 0.0209 | 0.0201 |
|  | hyp(PPT_year_): slope 2 | $\alpha_{7}$ | -0.0069 | 0.0022 |
|  | hyp(PPT_year_): x position | $\alpha_{8}$ | 922.86 | 72.39 |
|  | Greco B | $\alpha_{9}$ | 17.29 | 3.60 |
|  | Greco C | $\alpha_{10}$ | 9.01 | 2.29 |
|  | Greco D | $\alpha_{11}$ | 9.32 | 2.28 |
|  | Greco J | $\alpha_{12}$ | -17.15 | 3.49 |
| f_2_ | RDI | $\beta_{1}$ | -1.108 | 0.321 |
|  | RDI | $\beta_{2}$ | -0.286 | 0.038 |
| f_3_ | D_q_ | $\gamma_{1}$ | -0.114 | 0.009 |
|  | D_q_ | $\gamma_{2}$ | 0.088 | 0.010 |
|  | D_q_ | $\gamma_{3}$ | 152.5 | 15.7 |

##### f. Fraxinus excelsior

Function f_1_ for Fraxinus excelsior is given by:

$$f_{1}\left( X_{e} \right)=\alpha_{0}+\alpha_{1}.pH+\alpha_{2}.C/N+\alpha_{3}.\mathrm{Tmax}_{\mathrm{summer}}$$

**Table of parameters for Fraxinus excelsior:**

| **Function** | **Corresponding factor** | **Parameter** | **Estimate** | **Std. Error** |
| --- | --- | --- | --- | --- |
| f_1_ | Intercept | $\alpha_{0}$ | 377.62 | 151.69 |
|  | pH | $\alpha_{1}$ | -29.24 | 12.98 |
|  | C/N | $\alpha_{2}$ | -4.03 | 1.75 |
|  | Tmax_summer_ | $\alpha_{3}$ | -3.40 | 1.79 |
| f_2_ | RDI | $\beta_{1}$ | -6.988 | 2.501 |
|  | RDI | $\beta_{2}$ | 0.080 | 0.079 |
| f_3_ | D_q_ | $\gamma_{1}$ | -0.200 | 0.039 |
|  | D_q_ | $\gamma_{2}$ | 0.074 | 0.025 |
|  | D_q_ | $\gamma_{3}$ | 77.6 | 6.1 |

##### g. Larix decidua

Function f_1_ for Larix decidua is given by:

$$f_{1}\left( X_{e} \right)=\alpha_{0}+\alpha_{1}.C/{N+\alpha_{2}.SWHC}+\alpha_{3}.SGDD$$

**Table of parameters for Larix decidua:**

| **Function** | **Corresponding factor** | **Parameter** | **Estimate** | **Std. Error** |
| --- | --- | --- | --- | --- |
| f_1_ | Intercept | $\alpha_{0}$ | 86.48 | 44.49 |
|  | C/N | $\alpha_{1}$ | -2.02 | 1.03 |
|  | SWHC | $\alpha_{2}$ | 0.40 | 0.20 |
|  | SGDD | $\alpha_{3}$ | 0.0140 | 0.0085 |
| f_2_ | RDI | $\beta_{1}$ | -2.694 | 1.019 |
|  | RDI | $\beta_{2}$ | -0.109 | 0.117 |
| f_3_ | D_q_ | $\gamma_{1}$ | -0.180 | 0.032 |
|  | D_q_ | $\gamma_{2}$ | 0.037 | 0.015 |
|  | D_q_ | $\gamma_{3}$ | 133.2 | 50.4 |

##### h. Picea abies

Function f_1_ for Picea abies is given by:

$$f_{1}\left( X_{e} \right)=\alpha_{0}+\alpha_{1}.\left( pH-\alpha_{2} \right)^{2}+\alpha_{3}.C/N+\alpha_{4}.SWHC+\alpha_{5}.RockOutcrops+hyp\left( \mathrm{Tmin}_{\mathrm{winter}}, p=\alpha_{6}, q=0, x_{0}=\alpha_{7} \right)+\alpha_{8}.\mathrm{Greco}_{C}+\alpha_{9}.\mathrm{Greco}_{H}$$

**Table of parameters for Picea abies:**

| **Function** | **Corresponding factor** | **Parameter** | **Estimate** | **Std. Error** |
| --- | --- | --- | --- | --- |
| f_1_ | Intercept | $\alpha_{0}$ | 100.11 | 20.63 |
|  | pH | $\alpha_{1}$ | -2.92 | 1.20 |
|  | pH | $\alpha_{2}$ | 4.35 | 0.41 |
|  | C/N | $\alpha_{3}$ | -1.53 | 0.40 |
|  | SWHC | $\alpha_{4}$ | 0.107 | 0.038 |
|  | RockOutcrops | $\alpha_{5}$ | -1.97 | 0.68 |
|  | hyp(Tmin_winter_): slope 1 | $\alpha_{6}$ | 7.49 | 3.62 |
|  | hyp( Tmin_winter_ ): x position | $\alpha_{7}$ | -3.39 | 0.57 |
|  | Greco C | $\alpha_{8}$ | -8.31 | 3.01 |
|  | Greco H | $\alpha_{9}$ | -8.88 | 3.90 |
| f_2_ | RDI | $\beta_{1}$ | -2.876 | 0.512 |
|  | RDI | $\beta_{2}$ | -0.050 | 0.056 |
| f_3_ | D_q_ | $\gamma_{1}$ | -0.152 | 0.020 |
|  | D_q_ | $\gamma_{2}$ | 0.143 | 0.026 |
|  | D_q_ | $\gamma_{3}$ | 109.468 | 19.586 |

##### i. Pinus halepensis

Function f_1_ for Pinus halepensis is given by:

$$f_{1}\left( X_{e} \right)=\alpha_{0}+\alpha_{1}.\mathrm{Tmin}_{\mathrm{winter}}$$

**Table of parameters for Pinus halepensis:**

| **Function** | **Corresponding factor** | **Parameter** | **Estimate** | **Std. Error** |
| --- | --- | --- | --- | --- |
| f_1_ | Intercept | $\alpha_{0}$ | 36.27 | 21.59 |
|  | Tmin_winter_ | $\alpha_{1}$ | 1.94 | 1.72 |
| f_2_ | RDI | $\beta_{1}$ | -0.819 | 0.804 |
|  | RDI | $\beta_{2}$ | -0.270 | 0.111 |
| f_3_ | D_q_ | $\gamma_{1}$ | -0.217 | 0.067 |
|  | D_q_ | $\gamma_{2}$ | 0.106 | 0.062 |
|  | D_q_ | $\gamma_{3}$ | 98.3 | 30.9 |

##### j. Pinus nigra laricio

Function f_1_ for Pinus nigra laricio is given by:

$$f_{1}\left( X_{e} \right)=\alpha_{0}+\alpha_{1}.C/{N+\alpha_{2}.RockOutcrops}+\alpha_{3}.Limestone$$

**Table of parameters for Pinus nigra laricio:**

| **Function** | **Corresponding factor** | **Parameter** | **Estimate** | **Std. Error** |
| --- | --- | --- | --- | --- |
| f_1_ | Intercept | $\alpha_{0}$ | 85.69 | 20.62 |
|  | C/N | $\alpha_{1}$ | -0.98 | 0.34 |
|  | RockOutcrops | $\alpha_{2}$ | -2.94 | 0.98 |
|  | Limestone | $\alpha_{3}$ | -15.40 | 5.16 |
| f_2_ | RDI | $\beta_{1}$ | -3.389 | 0.868 |
|  | RDI | $\beta_{2}$ | 0.064 | 0.084 |
| f_3_ | D_q_ | $\gamma_{1}$ | -0.162 | 0.024 |
|  | D_q_ | $\gamma_{2}$ | 0.110 | 0.022 |
|  | D_q_ | $\gamma_{3}$ | 99.9 | 14.8 |

##### k. Pinus nigra nigra

Function f_1_ for Pinus nigra nigra is given by:

$$f_{1}\left( X_{e} \right)=\alpha_{0}+\alpha_{1}.C/{N+\alpha_{2}.\mathrm{Tmax}_{\mathrm{summer}}}+\alpha_{3}.\mathrm{Greco}_{F}$$

**Table of parameters for Pinus nigra nigra:**

| **Function** | **Corresponding factor** | **Parameter** | **Estimate** | **Std. Error** |
| --- | --- | --- | --- | --- |
| f_1_ | Intercept | $\alpha_{0}$ | 377.64 | 152.72 |
|  | C/N | $\alpha_{1}$ | -3.63 | 1.51 |
|  | Tmax_summer_ | $\alpha_{2}$ | 47.62 | 25.77 |
|  | Greco F | $\alpha_{3}$ | -8.85 | 3.61 |
| f_2_ | RDI | $\beta_{1}$ | -1.132 | 0.688 |
|  | RDI | $\beta_{2}$ | -0.186 | 0.079 |
| f_3_ | D_q_ | $\gamma_{1}$ | -0.217 | 0.046 |
|  | D_q_ | $\gamma_{2}$ | 0.091 | 0.035 |
|  | D_q_ | $\gamma_{3}$ | 62.1 | 5.9 |

##### l. Pinus pinaster

Function f_1_ for Pinus pinaster is given by:

$$f_{1}\left( X_{e} \right)=\alpha_{0}+hyp\left( pH, p=0, q=\alpha_{1}, x_{0}=\alpha_{2} \right)+\alpha_{3}.C/N+\alpha_{4}.Limestone+\alpha_{5}.RockOutcrops+\alpha_{6}.\mathrm{Soil}_{\mathrm{SH}}+hyp\left( \mathrm{wd}_{\mathrm{summer}}, p=0, q=\alpha_{7}, x_{0}=\alpha_{8} \right)+\alpha_{9}.\mathrm{Greco}_{I}$$

**Table of parameters for Pinus pinaster:**

| **Function** | **Corresponding factor** | **Parameter** | **Estimate** | **Std. Error** |
| --- | --- | --- | --- | --- |
| f_1_ | Intercept | $\alpha_{0}$ | 182.14 | 17.74 |
|  | hyp(pH): slope 2 | $\alpha_{1}$ | -12.83 | 2.91 |
|  | hyp(pH): x position | $\alpha_{2}$ | 4.76 | 0.21 |
|  | C/N | $\alpha_{3}$ | -2.76 | 0.35 |
|  | Limestone | $\alpha_{4}$ | -18.07 | 7.96 |
|  | RockOutcrops | $\alpha_{5}$ | -2.87 | 0.68 |
|  | Soil_SH_ | $\alpha_{6}$ | 10.63 | 2.44 |
|  | hyp(wd_summer_): slope 2 | $\alpha_{7}$ | -0.23 | 0.05 |
|  | hyp(wd_summer_): x position | $\alpha_{8}$ | 241.37 | 9.85 |
|  | Greco I | $\alpha_{9}$ | 108.62 | 49.99 |
| f_2_ | RDI | $\beta_{1}$ | -1.342 | 0.252 |
|  | RDI | $\beta_{2}$ | -0.086 | 0.033 |
| f_3_ | D_q_ | $\gamma_{1}$ | -0.163 | 0.009 |
|  | D_q_ | $\gamma_{2}$ | 0.088 | 0.006 |
|  | D_q_ | $\gamma_{3}$ | 77.6 | 3.8 |

##### m. Pinus sylvestris

Function f_1_ for Pinus sylvestris is given by:

$$f_{1}\left( X_{e} \right)=\alpha_{0}+\alpha_{1}.pH+hyp\left( C/N, p=\alpha_{2}, q=0, x_{0}=\alpha_{3} \right)+\alpha_{4}.SWHC+\alpha_{5}.\mathrm{Tmax}_{\mathrm{summer}}+\alpha_{6}.\mathrm{Greco}_{H}+\alpha_{7}.\mathrm{Greco}_{A, B,C,D}$$

For Pinus sylvestris, Greco A, B, C, and D were grouped, corresponding to half north of France.

**Table of parameters for Pinus sylvestris:**

| **Function** | **Corresponding factor** | **Parameter** | **Estimate** | **Std. Error** |
| --- | --- | --- | --- | --- |
| f_1_ | Intercept | $\alpha_{0}$ | 236.25 | 46.25 |
|  | pH | $\alpha_{1}$ | -7.81 | 1.88 |
|  | hyp(C/N): slope 1 | $\alpha_{2}$ | -3.40 | 0.92 |
|  | hyp(C/N): x position | $\alpha_{3}$ | 26.07 | 0.84 |
|  | SWHC | $\alpha_{4}$ | 0.18 | 0.06 |
|  | Tmax_summer_ | $\alpha_{5}$ | -4.94 | 1.08 |
|  | Greco H | $\alpha_{6}$ | -16.83 | 4.64 |
|  | Greco A, B, C, or D | $\alpha_{7}$ | 22.27 | 6.48 |
| f_2_ | RDI | $\beta_{1}$ | -0.809 | 0.357 |
|  | RDI | $\beta_{2}$ | -0.263 | 0.051 |
| f_3_ | D_q_ | $\gamma_{1}$ | -0.210 | 0.018 |
|  | D_q_ | $\gamma_{2}$ | 0.056 | 0.010 |
|  | D_q_ | $\gamma_{3}$ | 72.5 | 4.0 |

##### n. Pseudotsuga menziesii

Function f_1_ for Pseudotsuga menziesii is given by:

$$f_{1}\left( X_{e} \right)=\alpha_{0}+\alpha_{1}.Limestone+\alpha_{2}.ExpoEW+\alpha_{3}.\mathrm{Tmax}_{\mathrm{summer}}$$

$$+\alpha_{4}.\mathrm{Greco}_{B}+\alpha_{5}.\mathrm{Greco}_{D}+\alpha_{6}.\mathrm{Greco}_{I}$$

**Table of parameters for Pseudotsuga menziesii:**

| **Function** | **Corresponding factor** | **Parameter** | **Estimate** | **Std. Error** |
| --- | --- | --- | --- | --- |
| f_1_ | Intercept | $\alpha_{0}$ | 109.31 | 20.46 |
|  | Limestone | $\alpha_{1}$ | -10.27 | 3.54 |
|  | ExpoEW | $\alpha_{2}$ | -2.61 | 1.01 |
|  | Tmax_summer_ | $\alpha_{3}$ | -2.39 | 0.60 |
|  | Greco B | $\alpha_{4}$ | -4.39 | 1.88 |
|  | Greco D | $\alpha_{5}$ | -7.05 | 3.03 |
|  | Greco I | $\alpha_{6}$ | -17.74 | 4.98 |
| f_2_ | RDI | $\beta_{1}$ | -3.409 | 0.453 |
|  | RDI | $\beta_{2}$ | 0.082 | 0.045 |
| f_3_ | D_q_ | $\gamma_{1}$ | -0.160 | 0.022 |
|  | D_q_ | $\gamma_{2}$ | 0.240 | 0.038 |
|  | D_q_ | $\gamma_{3}$ | 130.7 | 21.2 |

##### o. Quercus petraea

Function f_1_ for Quercus petraea is given by:

$$f_{1}\left( X_{e} \right)=\alpha_{0}+\alpha_{1}.C/N+\alpha_{2}.SWHC+\alpha_{3}.RockOutcrops+\alpha_{4}.Limestone+\alpha_{5}.\mathrm{Tmin}_{\mathrm{winter}}+hyp\left( \mathrm{Tmax}_{\mathrm{summer}}, p=0, q=\alpha_{6}, x_{0}=\alpha_{7} \right)+hyp\left( \mathrm{PPT}_{\mathrm{year}}, p=\alpha_{8}, q=0, x_{0}=\alpha_{9} \right)+$$

**Table of parameters for Quercus petraea:**

| **Function** | **Corresponding factor** | **Parameter** | **Estimate** | **Std. Error** |
| --- | --- | --- | --- | --- |
| f_1_ | Intercept | $\alpha_{0}$ | 111.59 | 17.45 |
|  | C/N | $\alpha_{1}$ | -1.81 | 0.33 |
|  | SWHC | $\alpha_{2}$ | 0.13 | 0.04 |
|  | RockOutcrops | $\alpha_{3}$ | -3.49 | 0.91 |
|  | Limestone | $\alpha_{4}$ | -25.32 | 5.01 |
|  | Tmin_winter_ | $\alpha_{5}$ | 8.19 | 1.66 |
|  | hyp(Tmax_summer_): slope 2 | $\alpha_{6}$ | -8.51 | 1.99 |
|  | hyp(Tmax_summer_): x position | $\alpha_{7}$ | 24.40 | 0.31 |
|  | hyp(PPT_year_): slope 1 | $\alpha_{8}$ | 0.12 | 0.05 |
|  | hyp(PPT_year_ ): x position | $\alpha_{9}$ | 750.19 | 25.21 |
| f_2_ | RDI | $\beta_{1}$ | -1.750 | 0.290 |
|  | RDI | $\beta_{2}$ | -0.072 | 0.035 |
| f_3_ | D_q_ | $\gamma_{1}$ | -0.216 | 0.014 |
|  | D_q_ | $\gamma_{2}$ | 0.050 | 0.007 |
|  | D_q_ | $\gamma_{3}$ | 105.7 | 4.2 |

##### p. Quercus pubescens

Function f_1_ for Quercus pubescens is given by:

$$f_{1}\left( X_{e} \right)=\alpha_{0}+\alpha_{1}.C/N+\alpha_{2}.SWHC$$

**Table of parameters for Quercus pubescens:**

| **Function** | **Corresponding factor** | **Parameter** | **Estimate** | **Std. Error** |
| --- | --- | --- | --- | --- |
| f_1_ | Intercept | $\alpha_{0}$ | 53.65 | 31.70 |
|  | C/N | $\alpha_{1}$ | -1.30 | 0.78 |
|  | SWHC | $\alpha_{2}$ | 0.17 | 0.11 |
| f_2_ | RDI | $\beta_{1}$ | -0.960 | 0.546 |
|  | RDI | $\beta_{2}$ | -0.192 | 0.080 |
| f_3_ | D_q_ | $\gamma_{1}$ | -0.263 | 0.061 |
|  | D_q_ | $\gamma_{2}$ | 0.084 | 0.051 |
|  | D_q_ | $\gamma_{3}$ | 84.6 | 7.8 |

##### q. Quercus robur

Function f_1_ for Quercus robur is given by:

$$f_{1}\left( X_{e} \right)=\alpha_{0}+\alpha_{1}.C/N+\alpha_{2}.SWHC+\alpha_{3}.Limestone+\alpha_{4}.\mathrm{Tmin}_{\mathrm{winter}}+hyp\left( \mathrm{wd}_{\mathrm{summer}}, p=0, q=\alpha_{5}, x_{0}=\alpha_{6} \right)$$

**Table of parameters for Quercus robur:**

| **Function** | **Corresponding factor** | **Parameter** | **Estimate** | **Std. Error** |
| --- | --- | --- | --- | --- |
| f_1_ | Intercept | $\alpha_{0}$ | 42.92 | 6.79 |
|  | C/N | $\alpha_{1}$ | -0.64 | 0.14 |
|  | SWHC | $\alpha_{2}$ | 0.065 | 0.026 |
|  | Limestone | $\alpha_{3}$ | -7.31 | 2.80 |
|  | Tmin_winter_ | $\alpha_{4}$ | 3.04 | 0.84 |
|  | hyp(wd_summer_): slope 2 | $\alpha_{5}$ | -0.13 | 0.06 |
|  | hyp(wd_summer_): x position | $\alpha_{6}$ | 207.81 | 14.78 |
| f_2_ | RDI | $\beta_{1}$ | -0.614 | 0.327 |
|  | RDI | $\beta_{2}$ | -0.309 | 0.044 |
| f_3_ | D_q_ | $\gamma_{1}$ | -0.142 | 0.012 |
|  | D_q_ | $\gamma_{2}$ | 0.091 | 0.012 |
|  | D_q_ | $\gamma_{3}$ | 130.2 | 9.5 |

##### r. Robinia pseudoacacia

Function f_1_ for Robinia pseudoacacia is given by:

$$f_{1}\left( X_{e} \right)=\alpha_{0}+\alpha_{1}.C/N+\alpha_{2}.SWHC+\alpha_{3}.SGDD$$

**Table of parameters for Robinia pseudoacacia:**

| **Function** | **Corresponding factor** | **Parameter** | **Estimate** | **Std. Error** |
| --- | --- | --- | --- | --- |
| f_1_ | Intercept | $\alpha_{0}$ | 158.58 | 136.71 |
|  | C/N | $\alpha_{1}$ | -11.51 | 9.36 |
|  | SWHC | $\alpha_{2}$ | 0.77 | 0.71 |
|  | SGDD | $\alpha_{3}$ | 0.03 | 0.04 |
| f_2_ | RDI | $\beta_{1}$ | -10.131 | 7.178 |
|  | RDI | $\beta_{2}$ | 0.194 | 0.192 |
| f_3_ | D_q_ | $\gamma_{1}$ | -0.336 | 0.103 |
|  | D_q_ | $\gamma_{2}$ | 0.033 | 0.024 |
|  | D_q_ | $\gamma_{3}$ | 85.5 | 27.0 |

#### 4. Models’ residuals

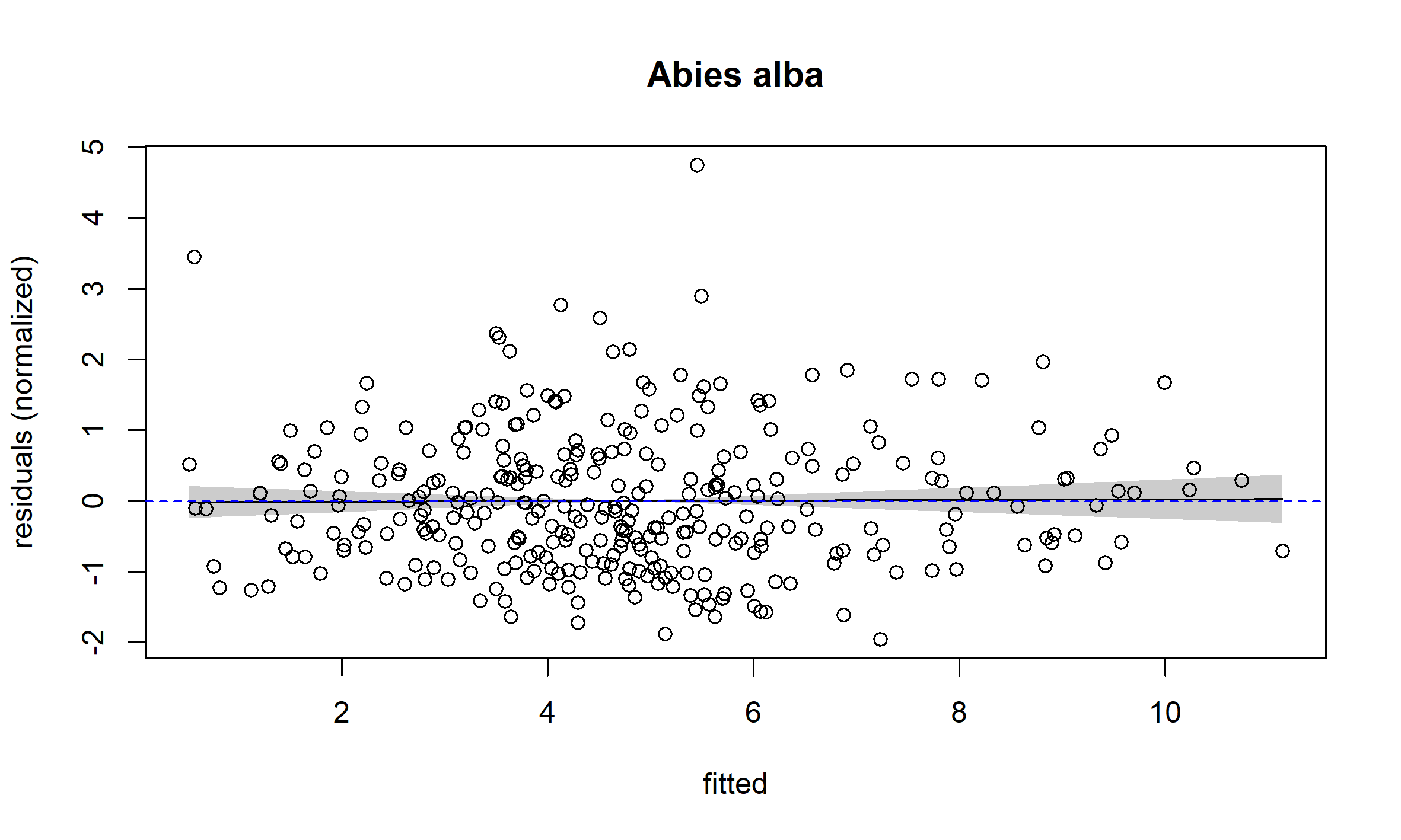

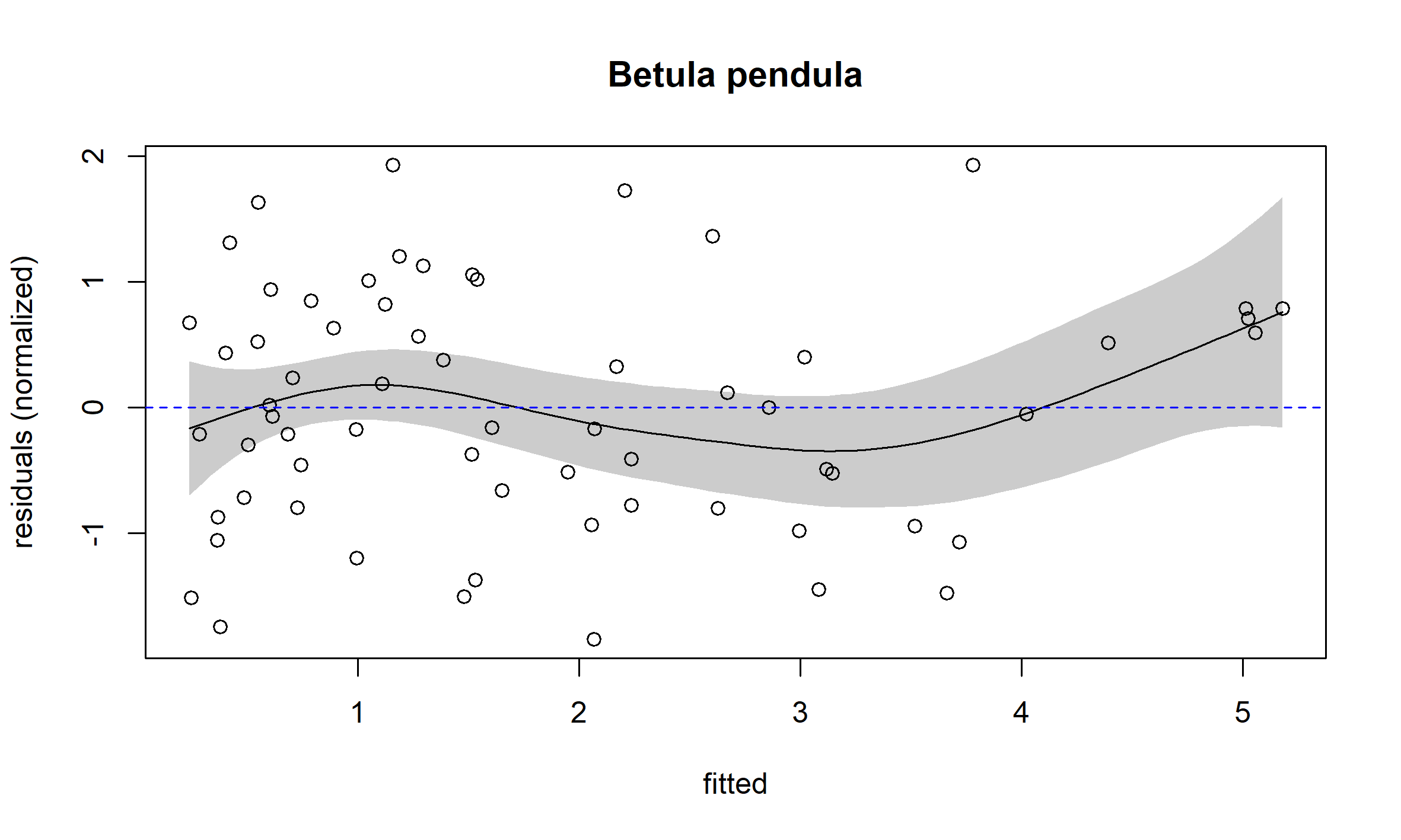

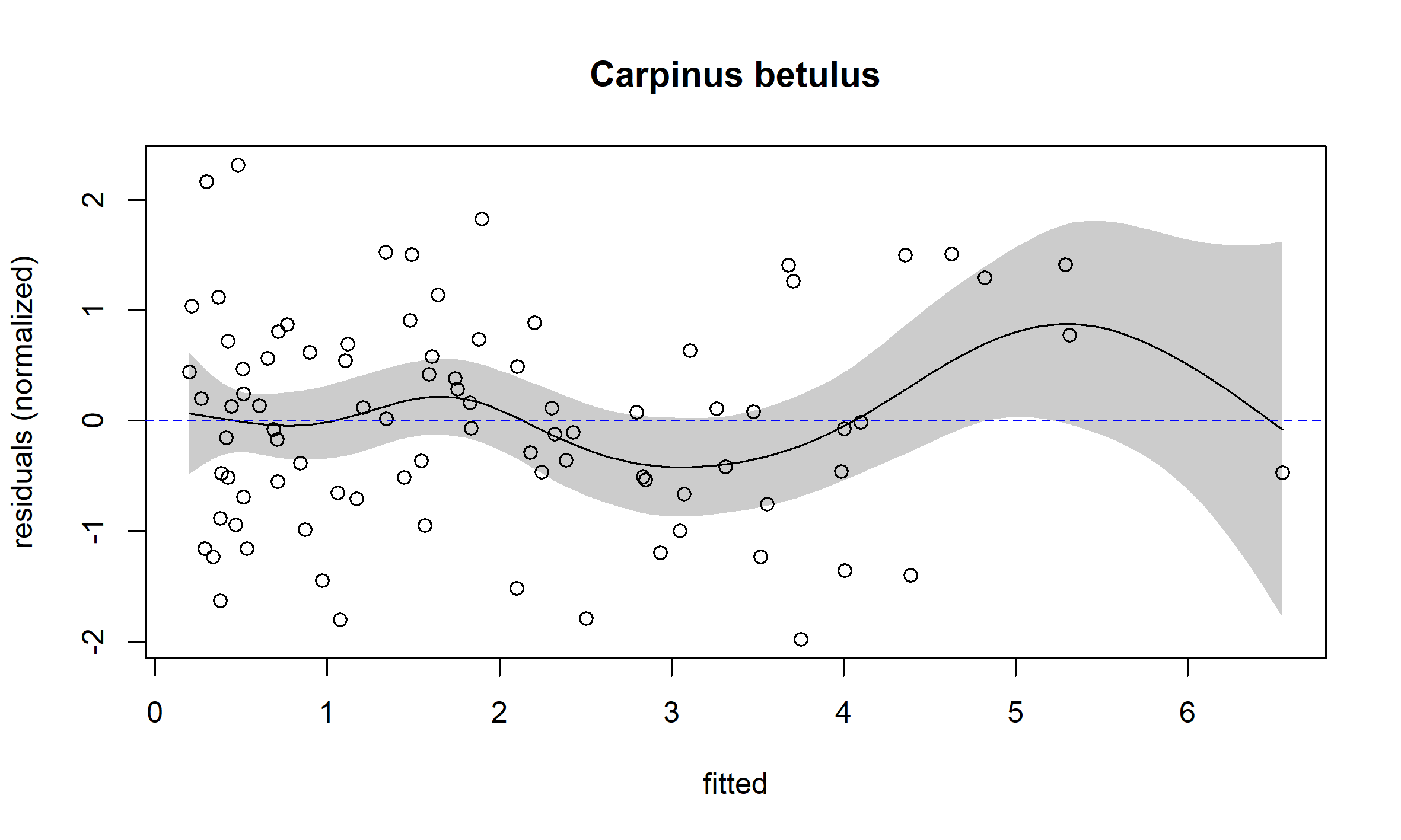

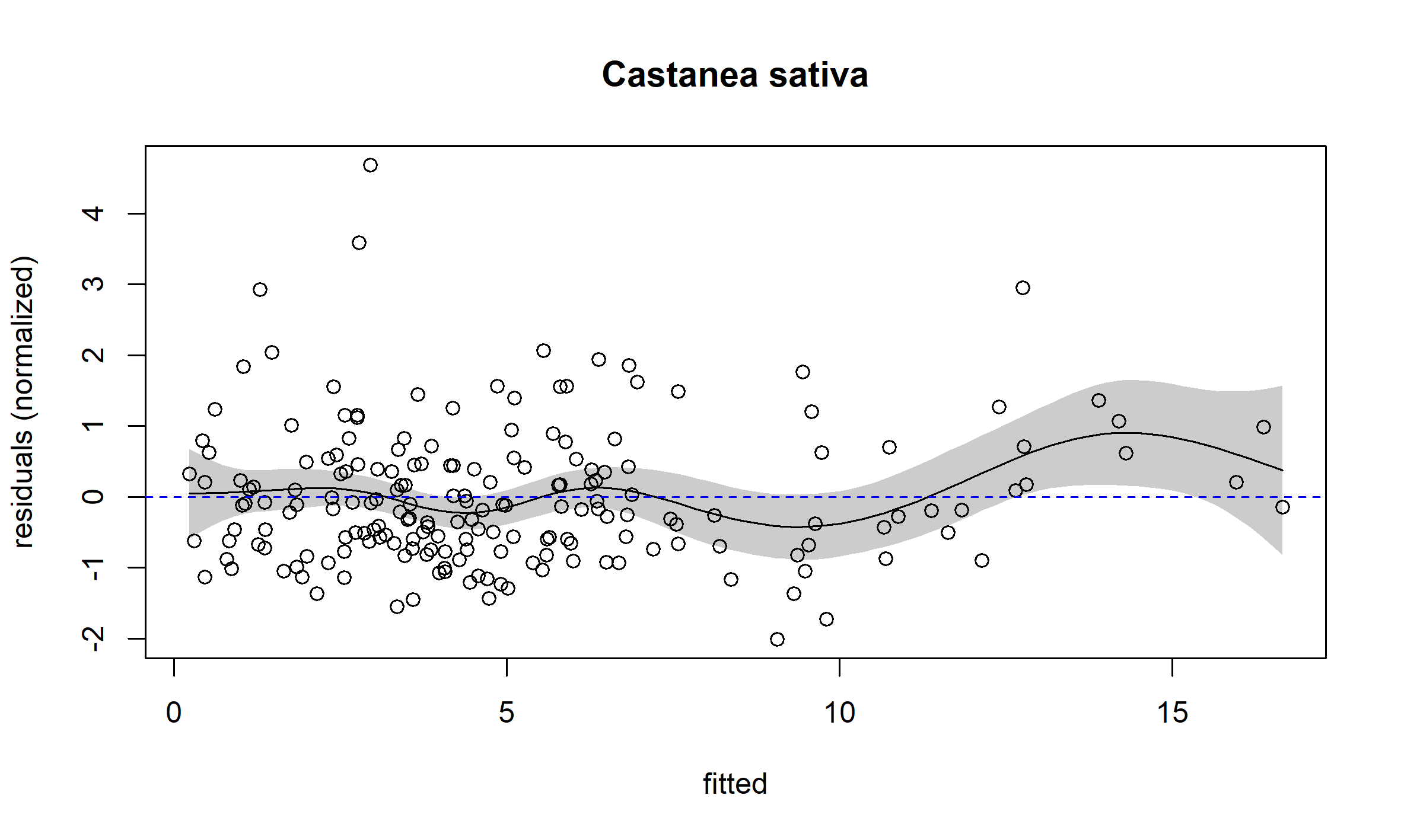

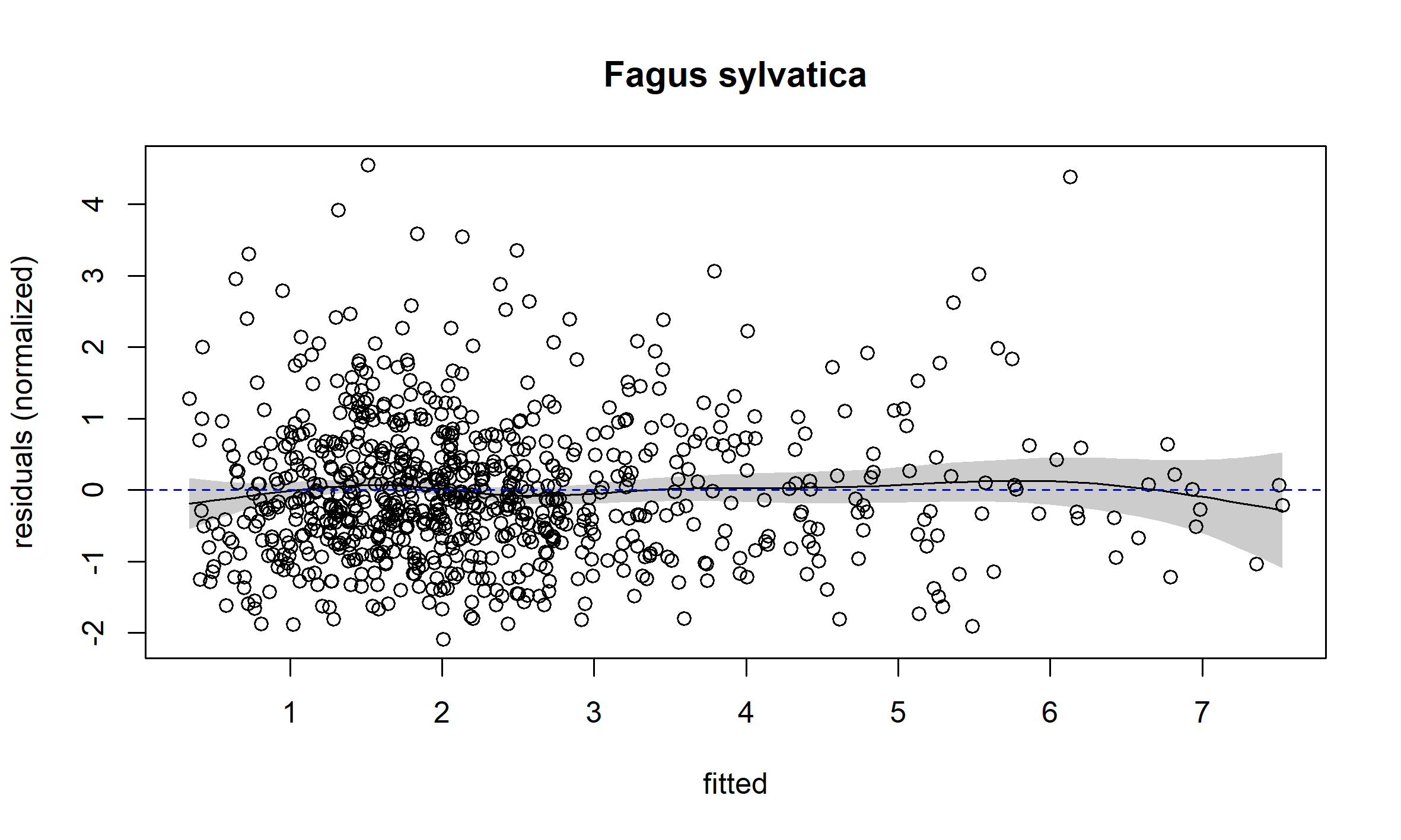

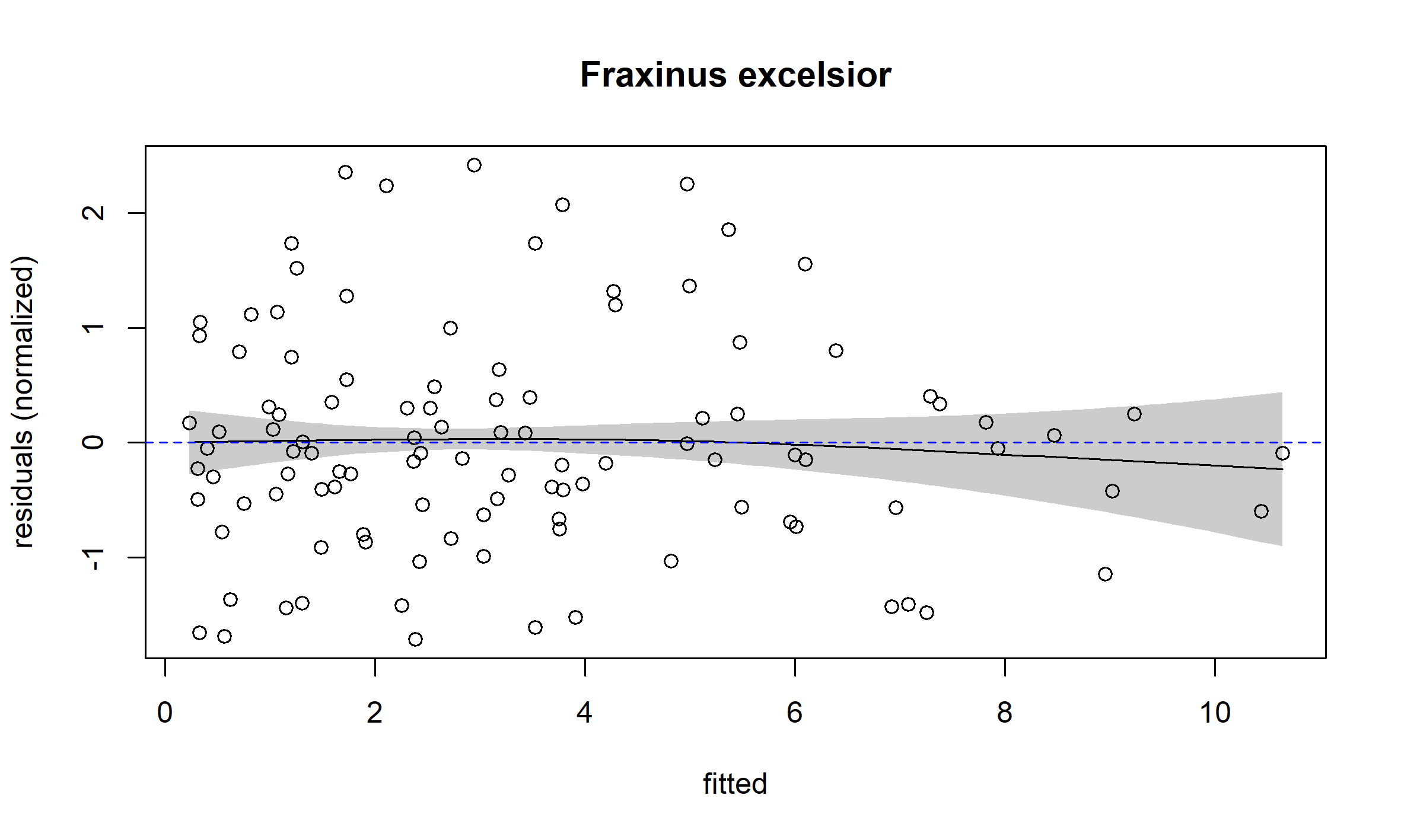

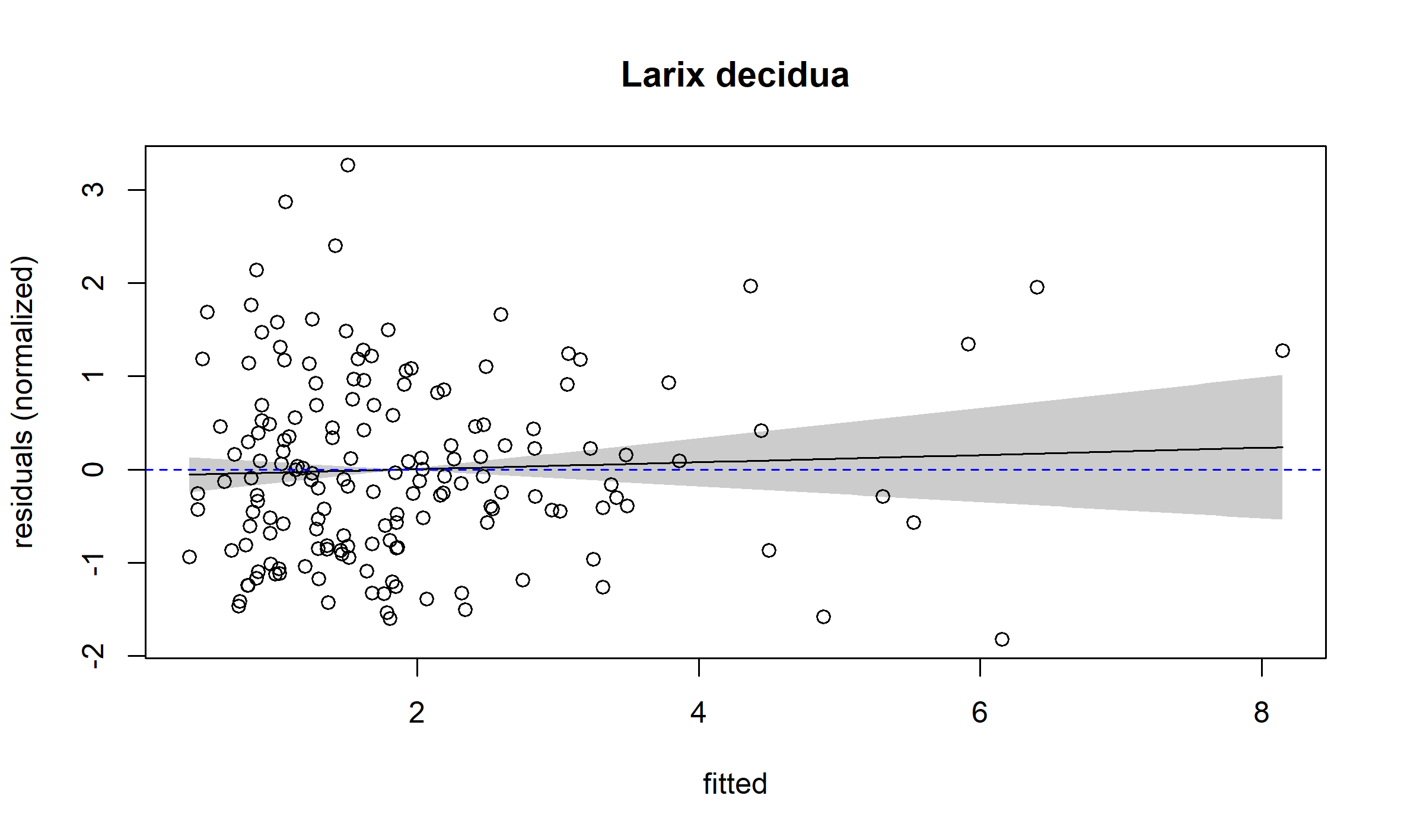

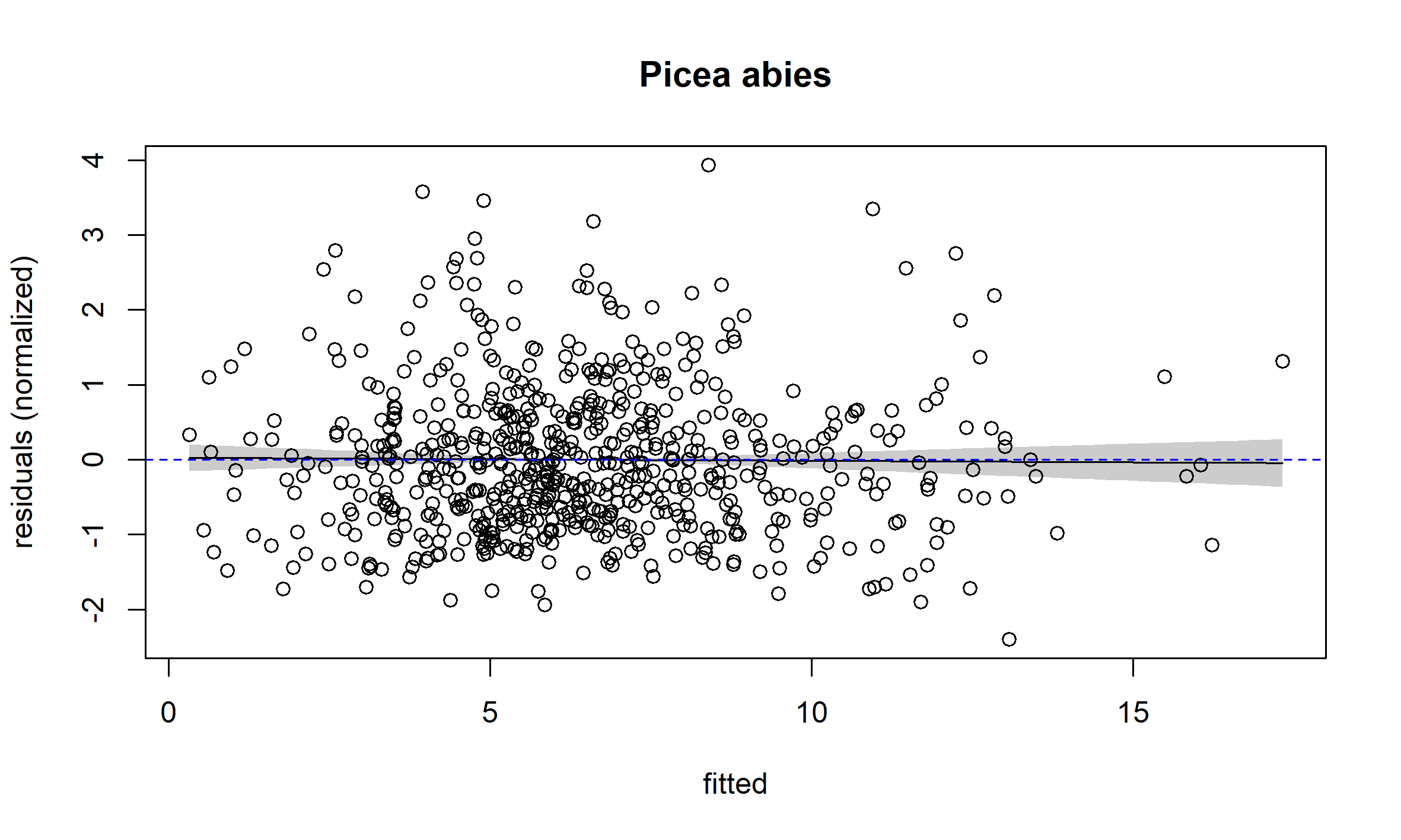

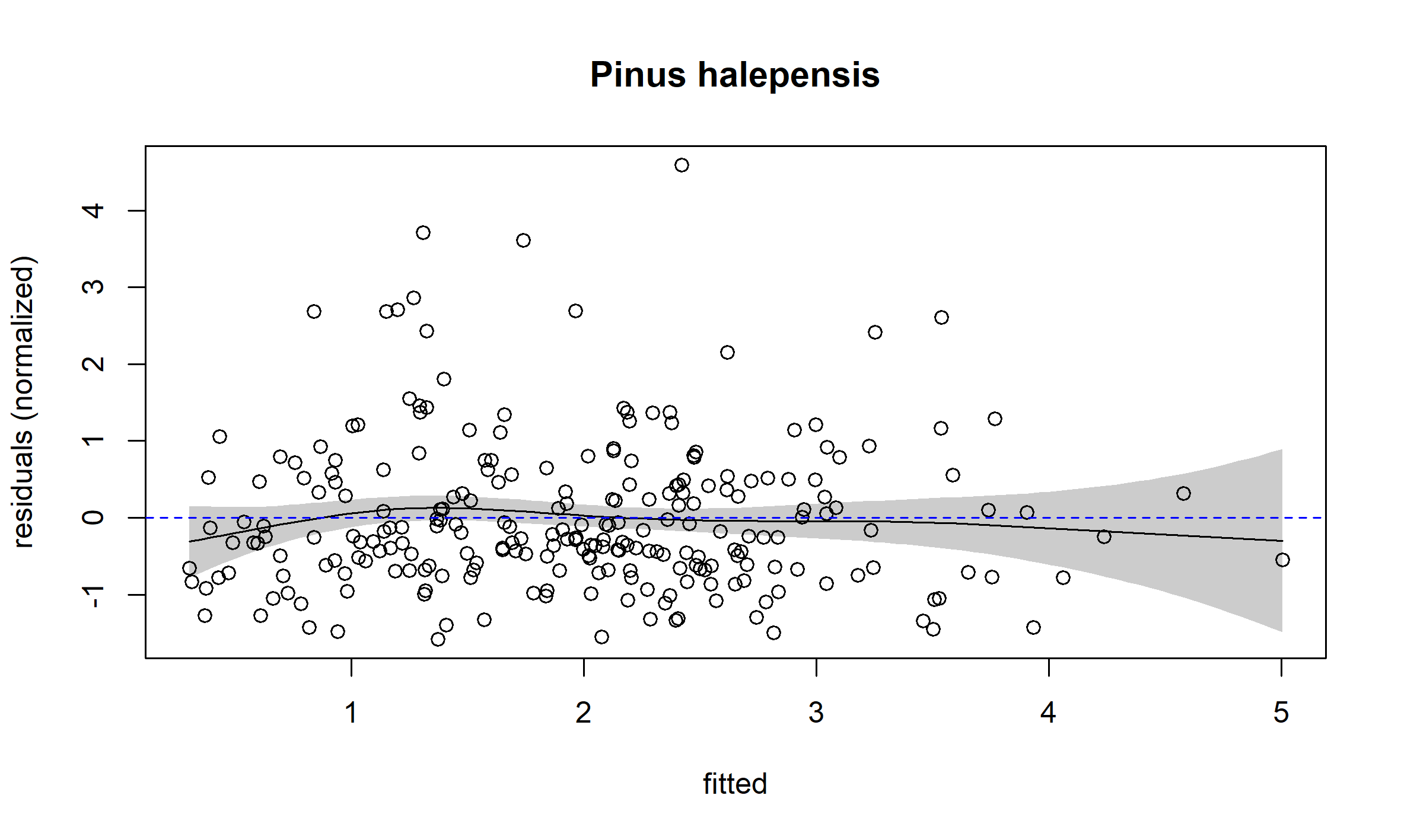

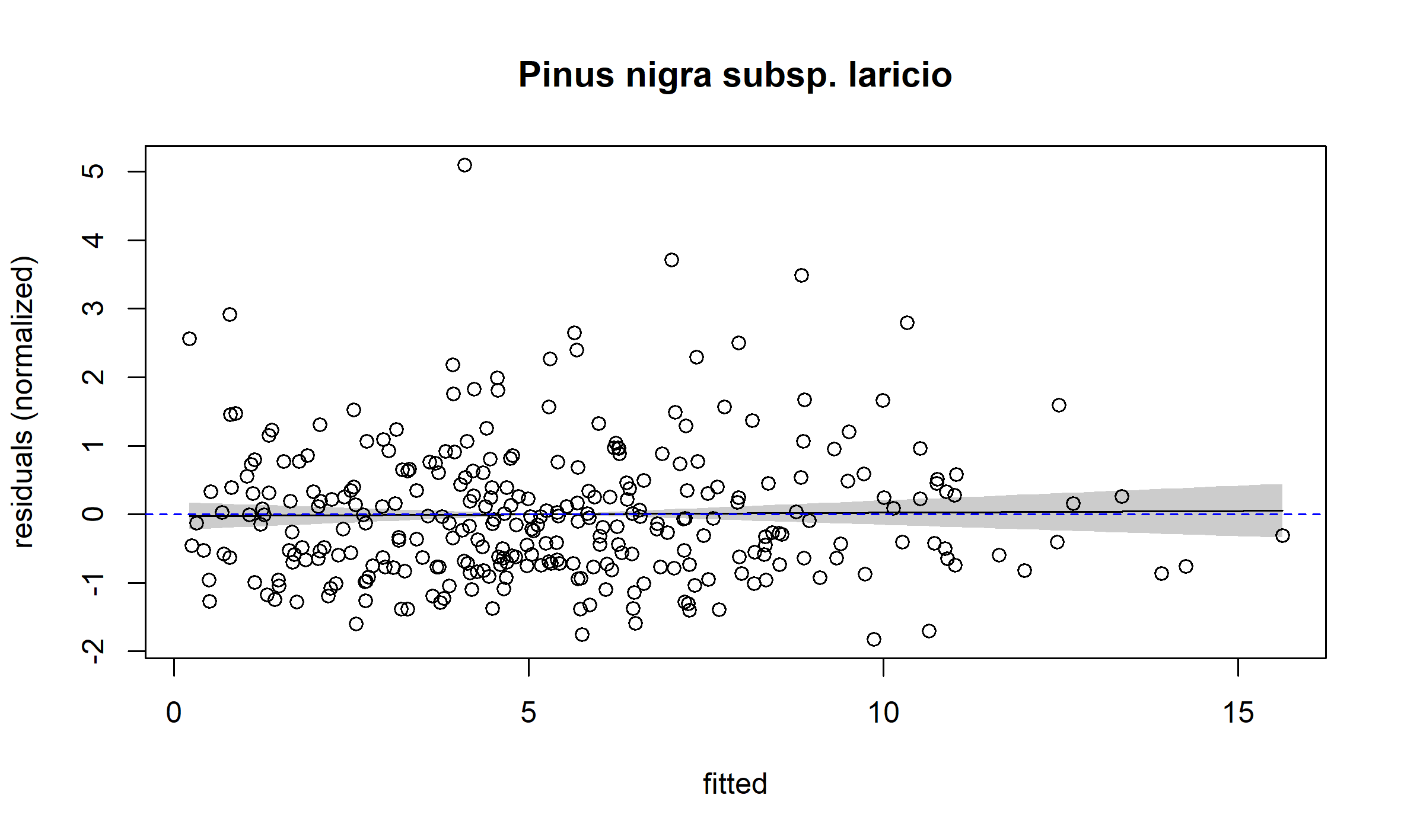

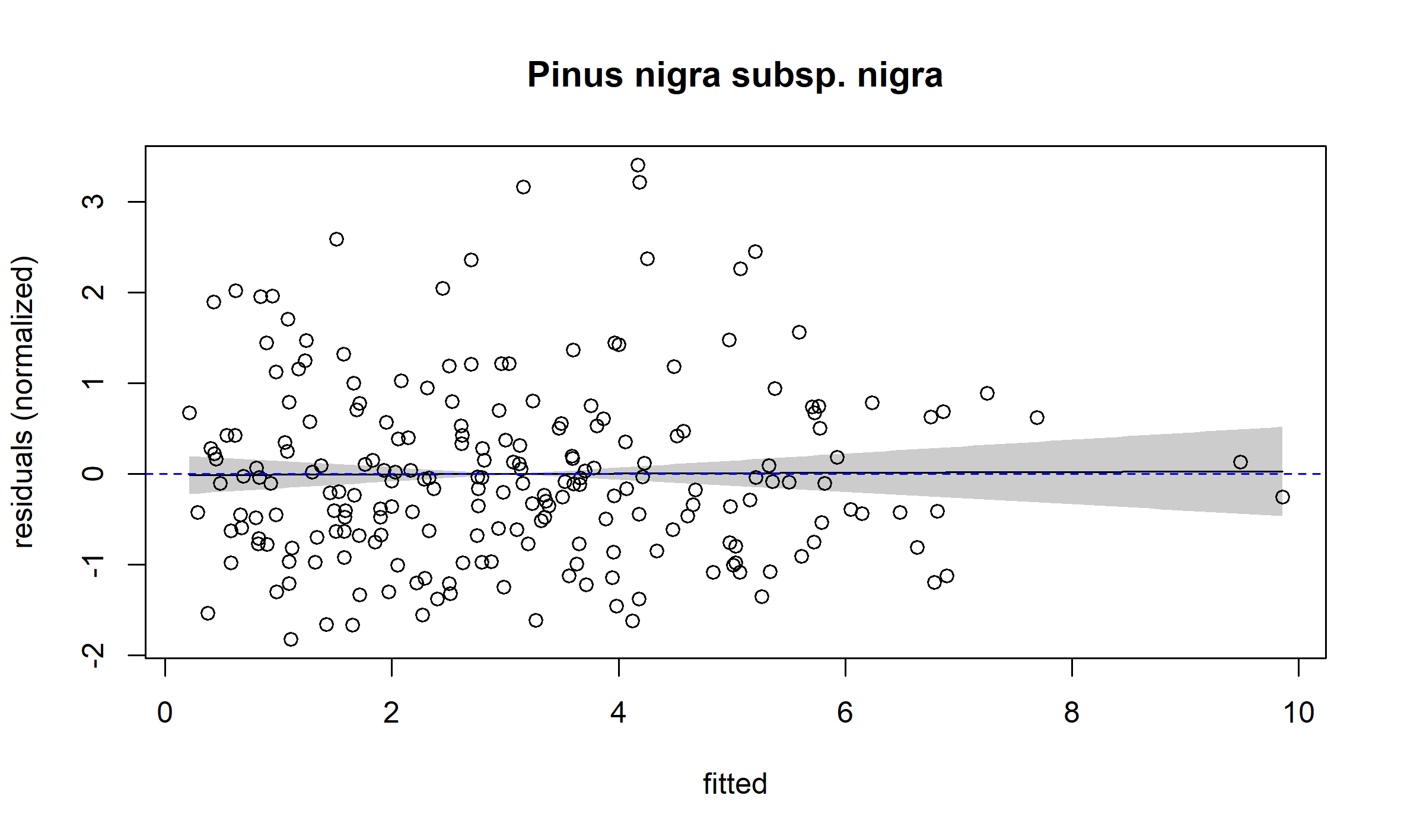

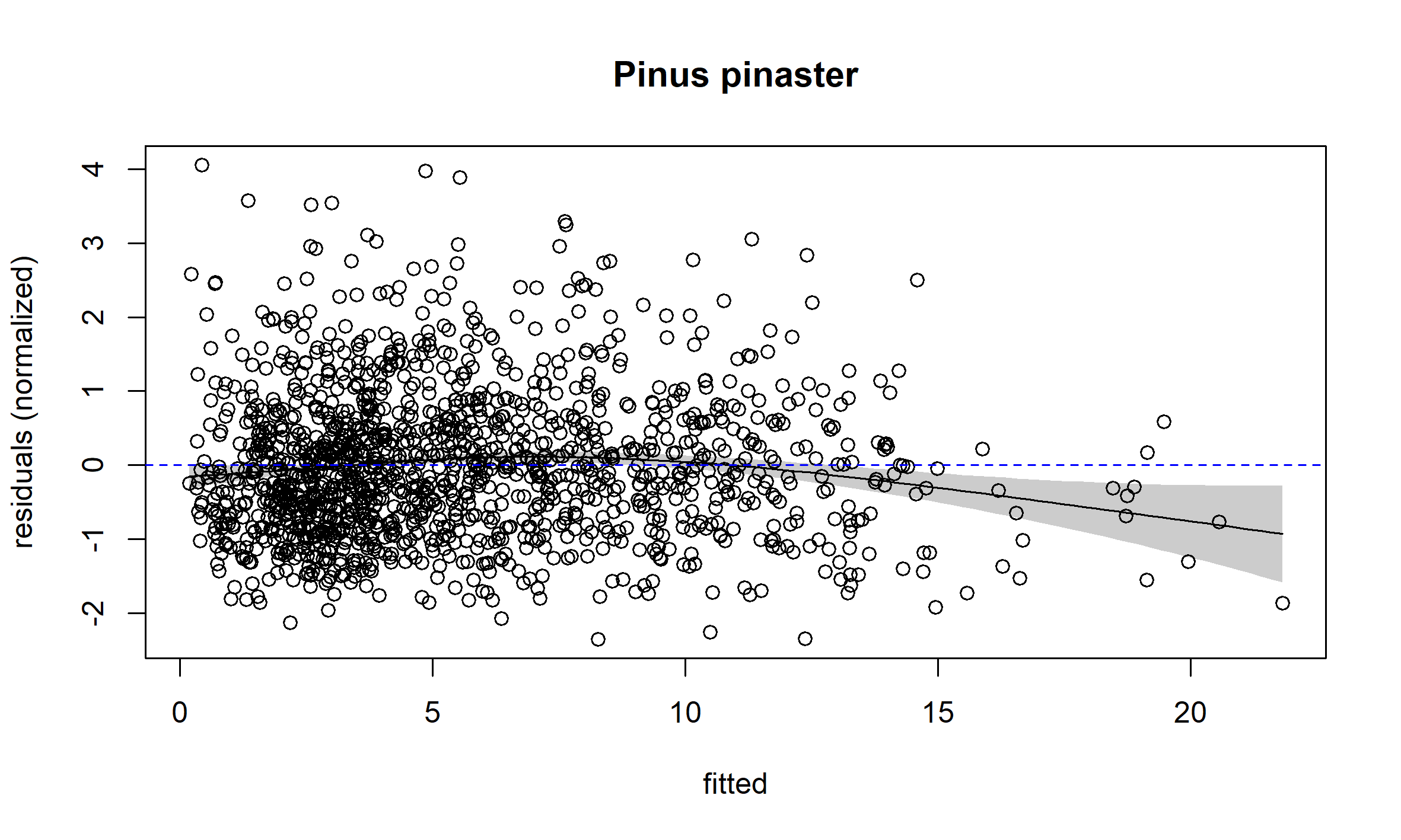

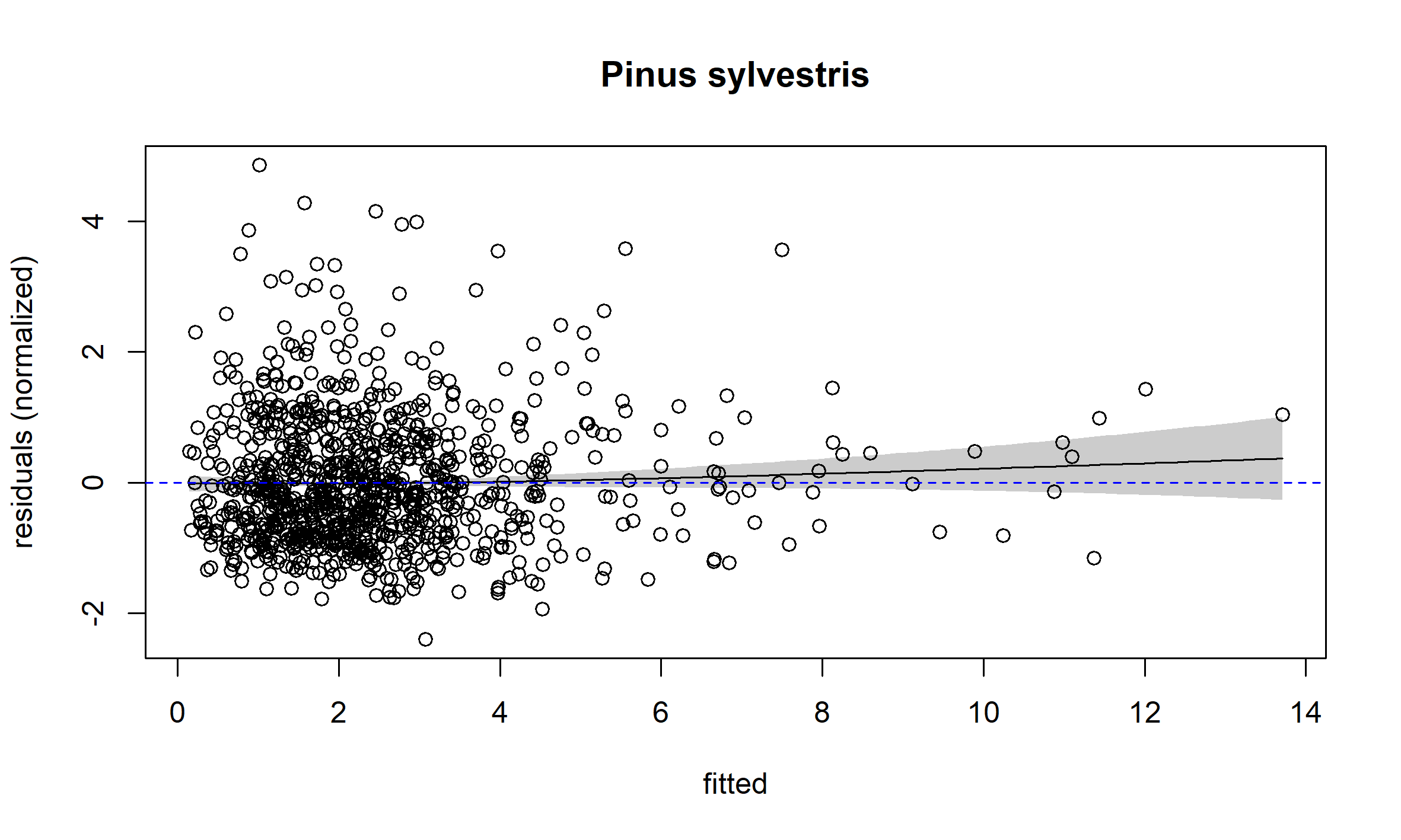

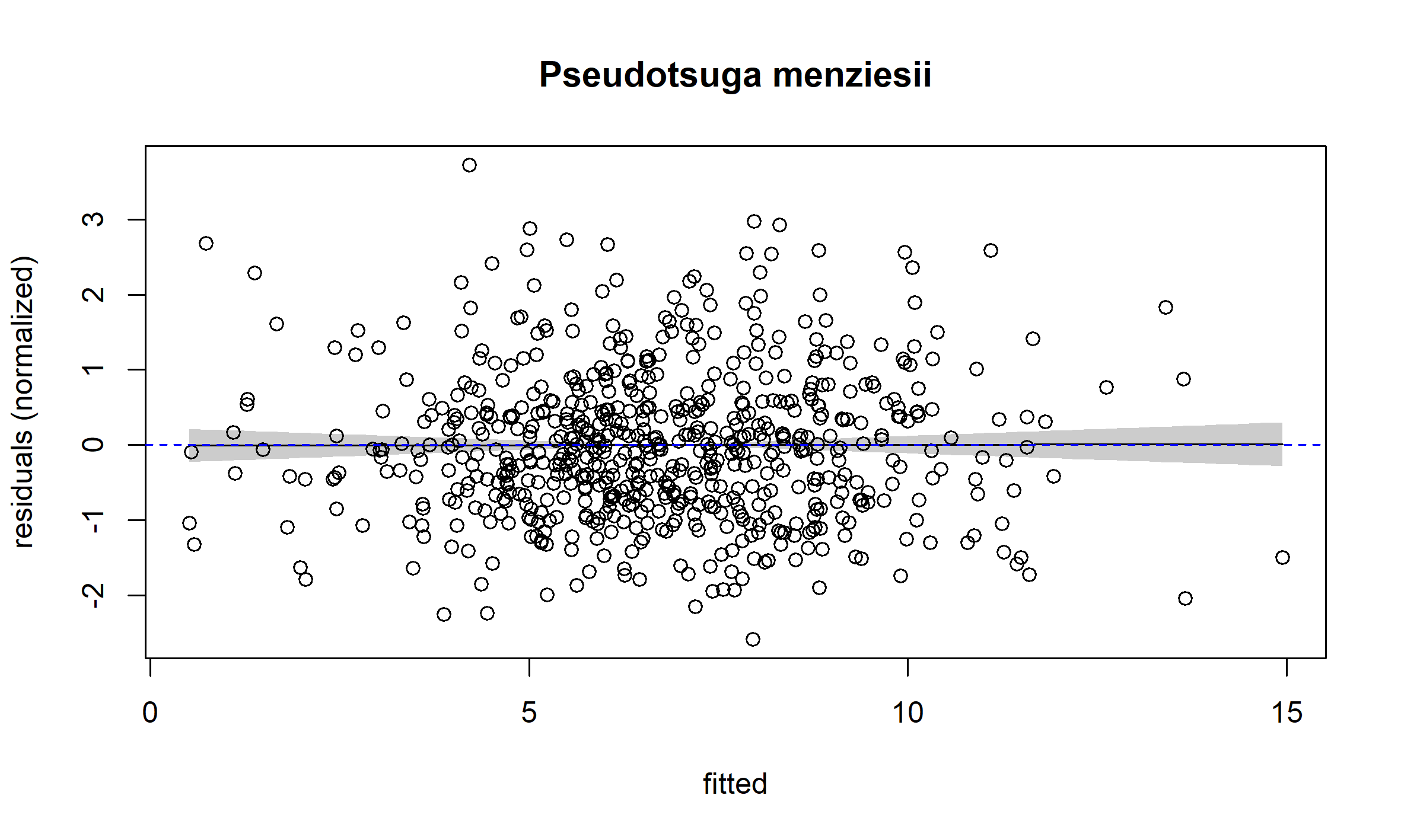

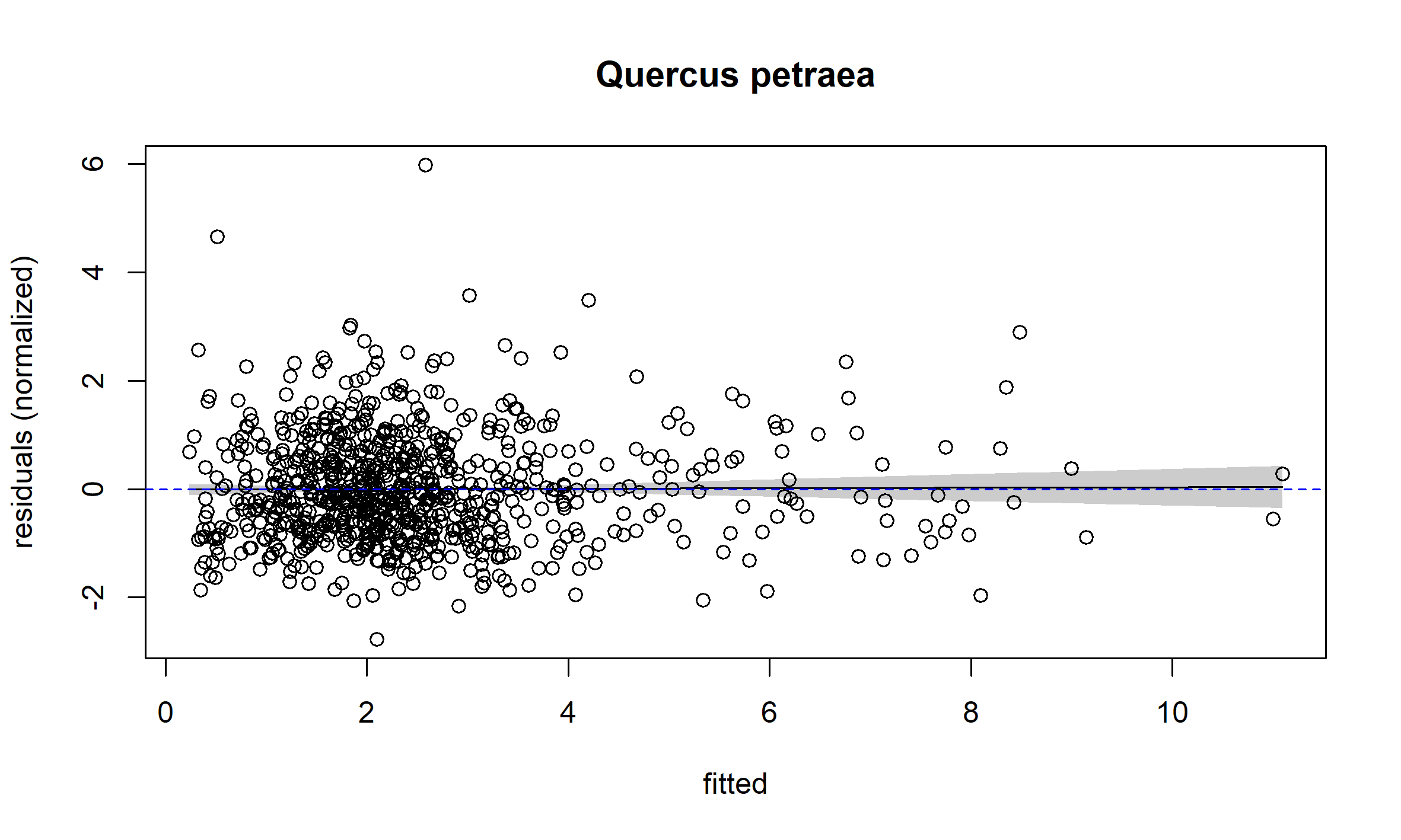

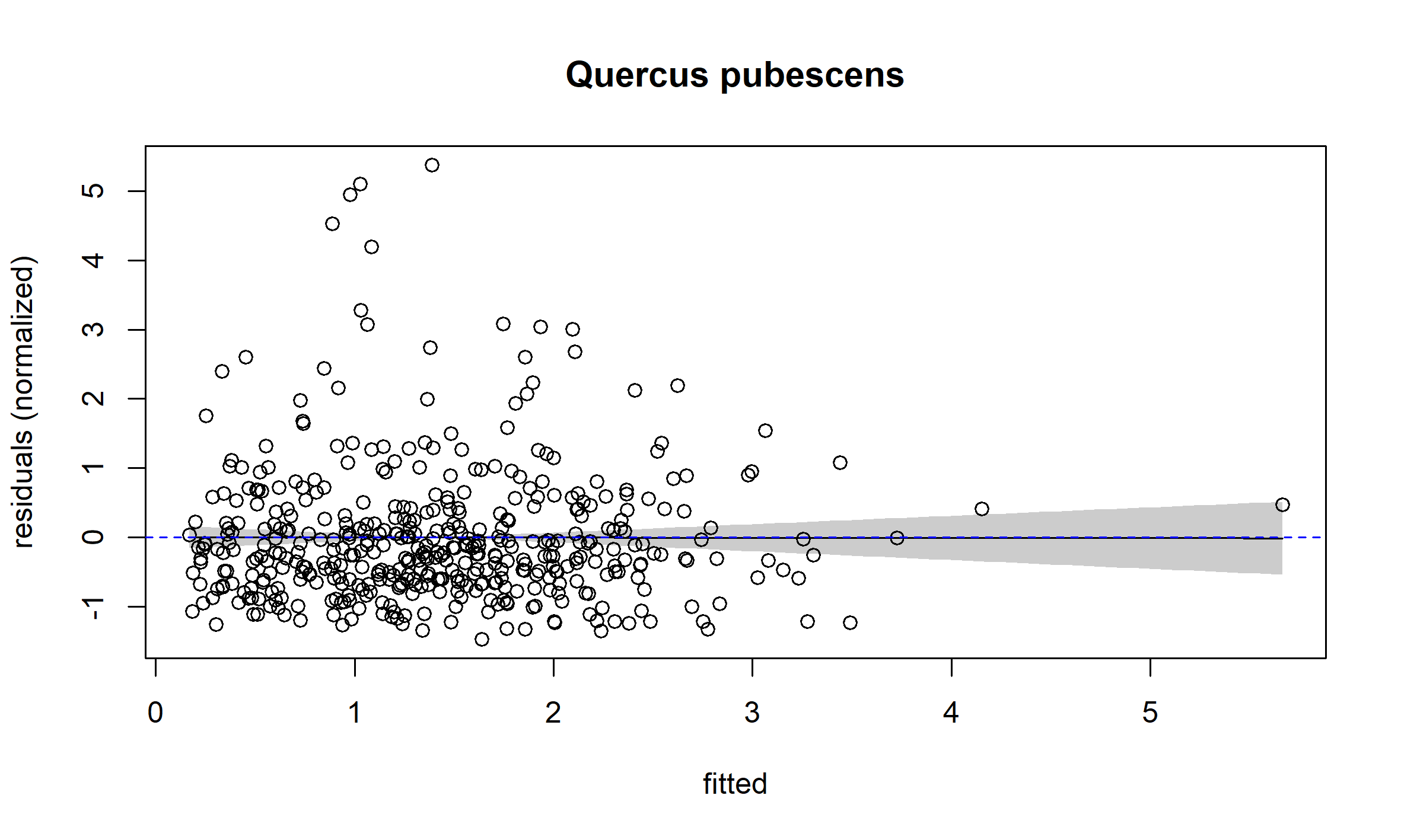

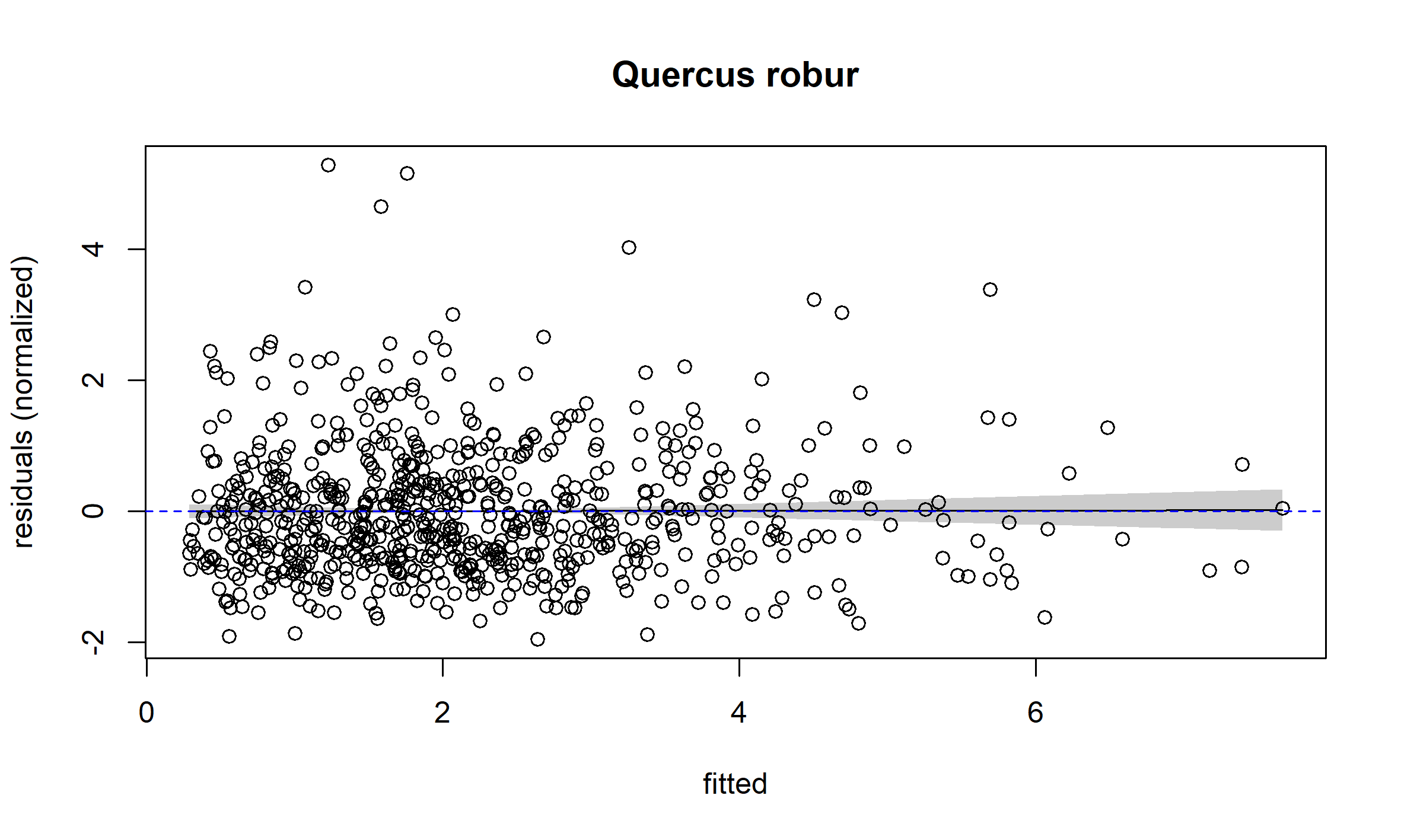

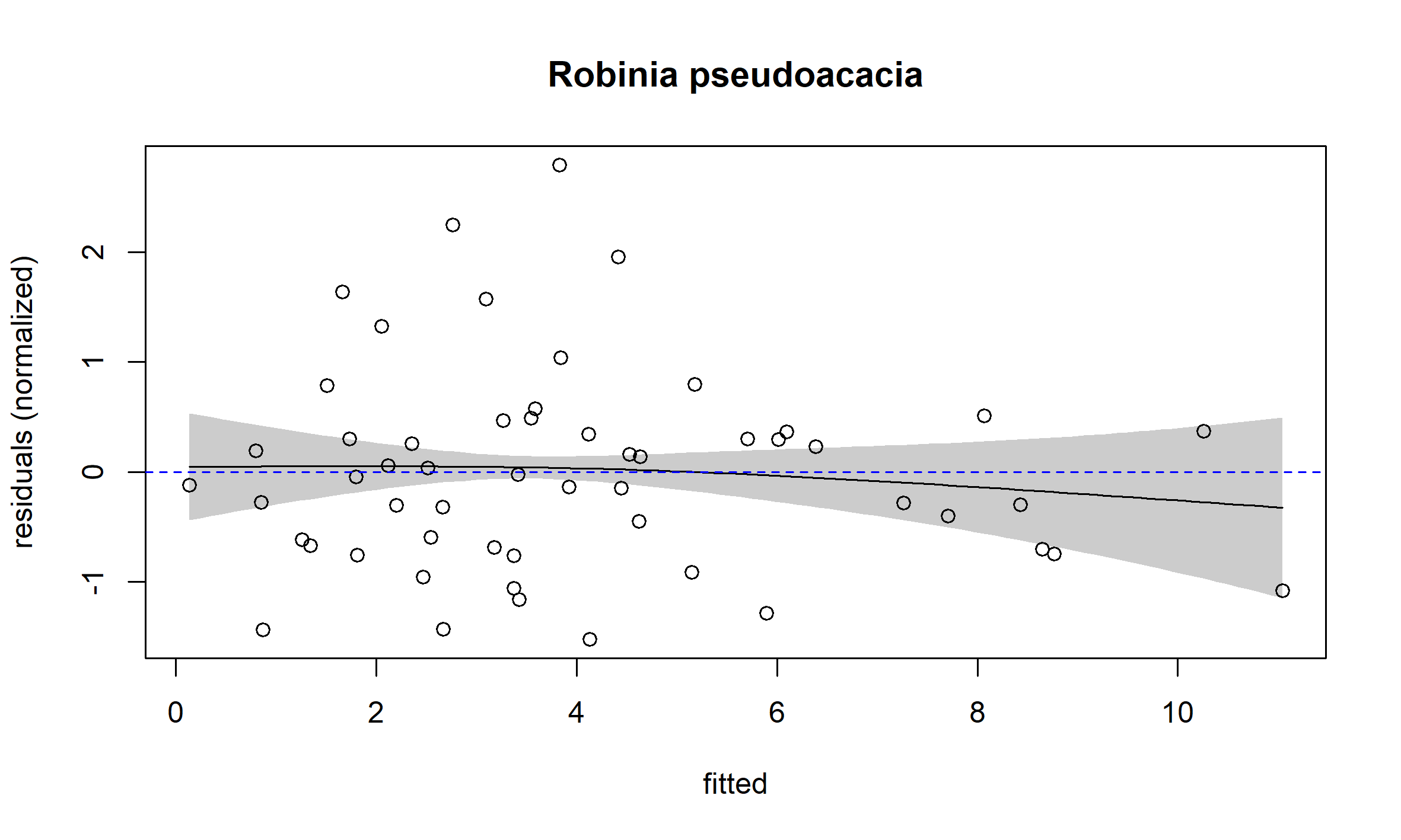

### D. Residuals of models without and with the hyperbolic forms of climate variable

In the calibration of our models, we identified a hyperbolic form for seven climate variables. For each of them, in the following sub-sections, we illustrate the interest of including the climate variable with a hyperbolic form.

For each of them, we calibrated a model similar for the final one, except that we excluded the climate variable. We then plotted the residuals of this model against the variable, and used a smoothing curve to assess the structure of the residuals (plots of the left side). We also plotted the residuals of the model including the climate variable – i.e., the final model – with the smoothing curve.

In all the seven cases, the hyperbolic form made it possible to avoid a structure in the residuals.

##### a. Abies alba – effect of variable SGDD

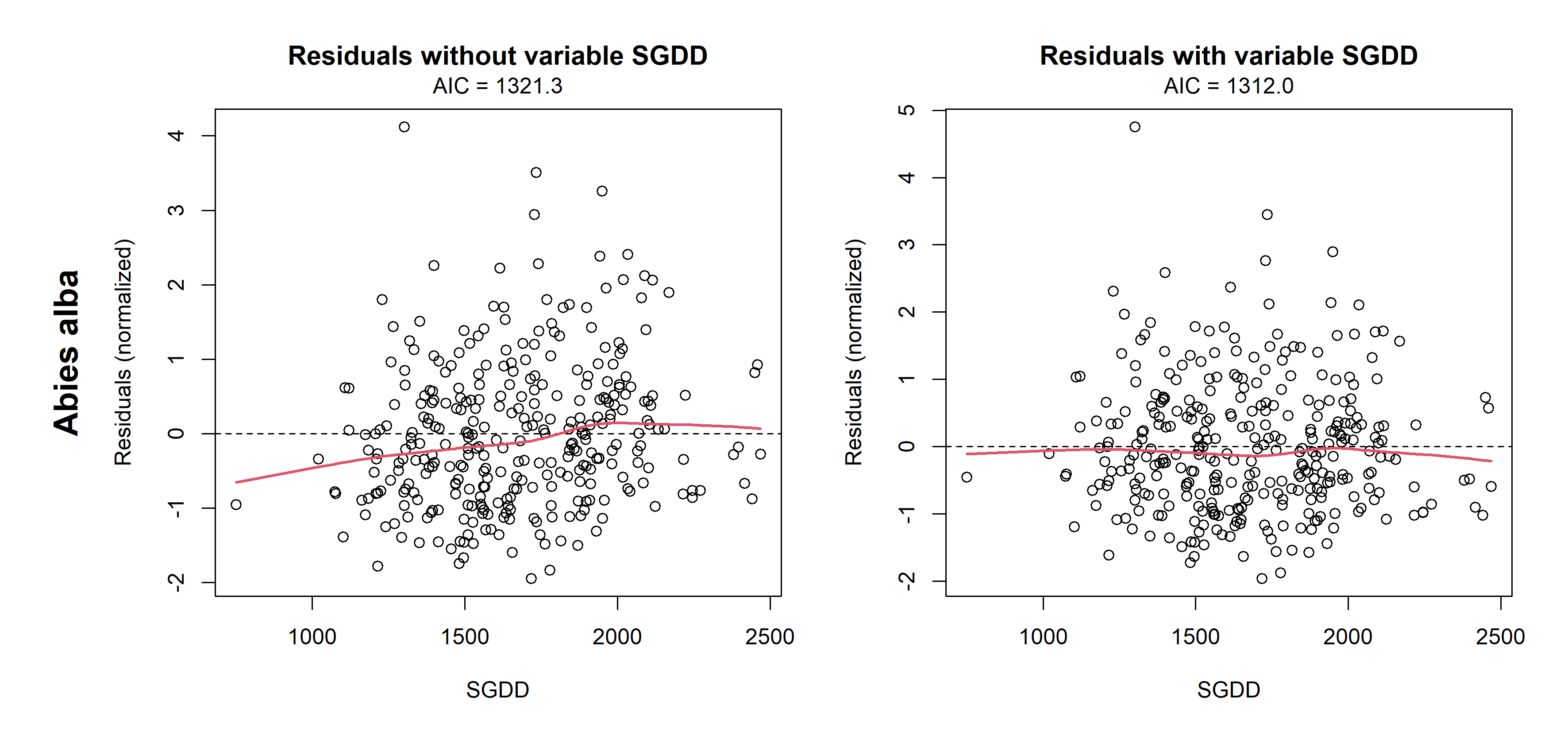

Figure D1. Models normalized residuals against SGDD for Abies alba.
On the left side, the residuals are for a model before the inclusion of the SGDD variable.
On the right side, the residuals are for the model with the inclusion of the SGDD variable.

##### b. Fagus sylvatica – effect of variable Annual Precipitation

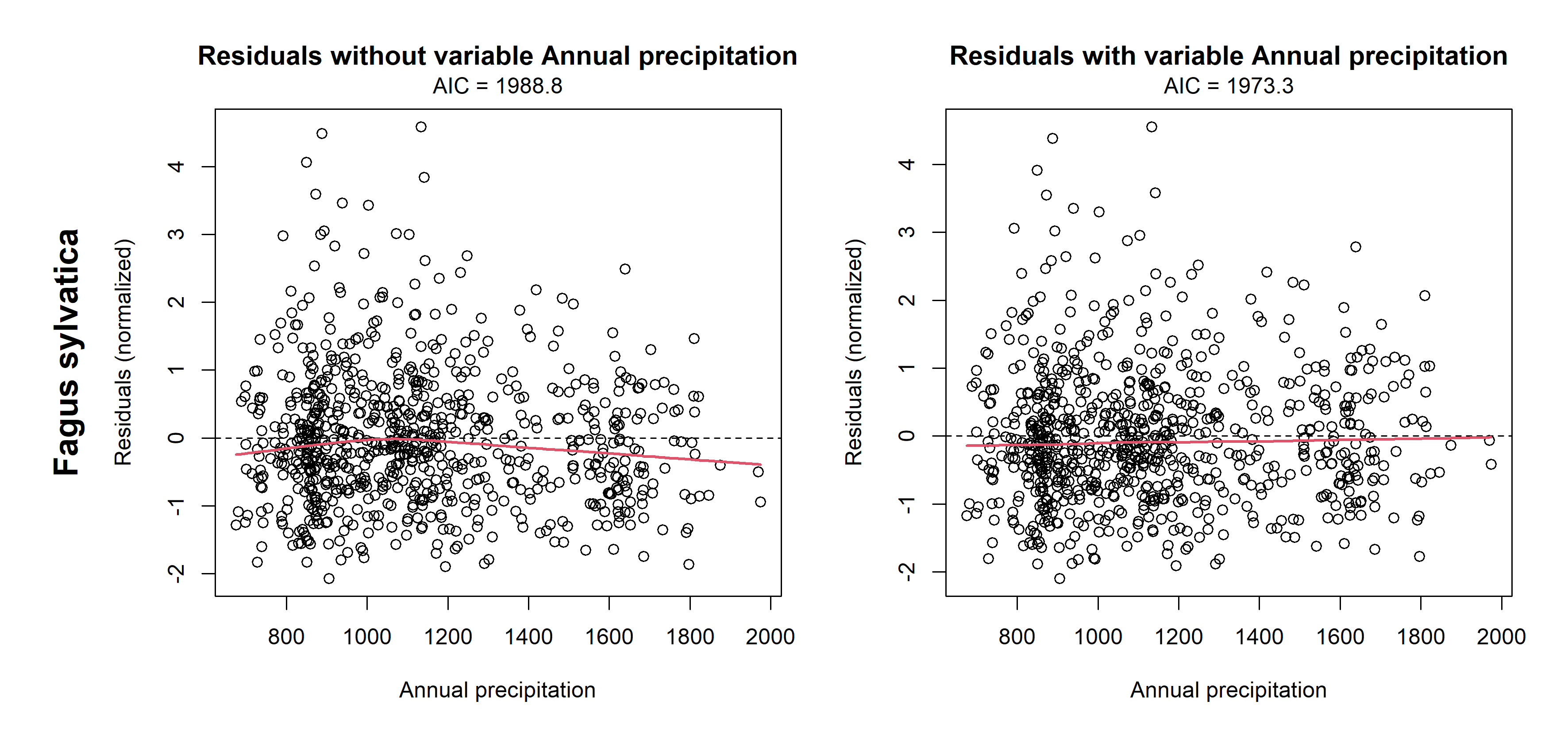

Figure D2. Models normalized residuals against Annual Precipitation for Fagus sylvatica.
On the left side, the residuals are for a model before the inclusion of the Annual precipitation variable.
On the right side, the residuals are for the model with the inclusion of the Annual precipitation variable.

##### c. Picea abies – effect of variable Winter Min Temperature

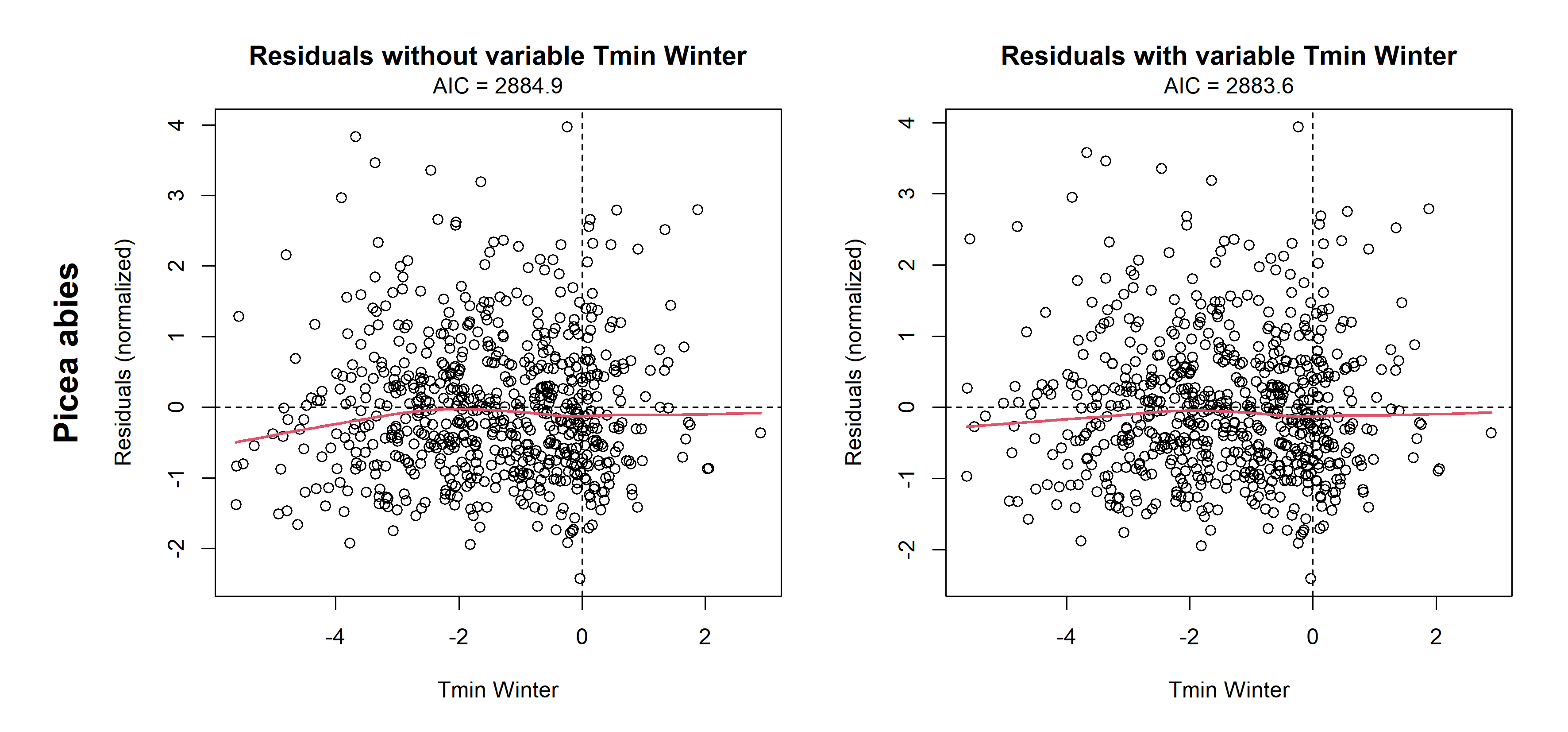

Figure D3. Models normalized residuals against Winter Min Temperature for Picea abies.
On the left side, the residuals are for a model before the inclusion of the Winter Min Temperature variable.
On the right side, the residuals are for the model with the inclusion of the Winter Min Temperature variable.

##### d. Pinus pinaster – effect of variable Summer Water Deficit

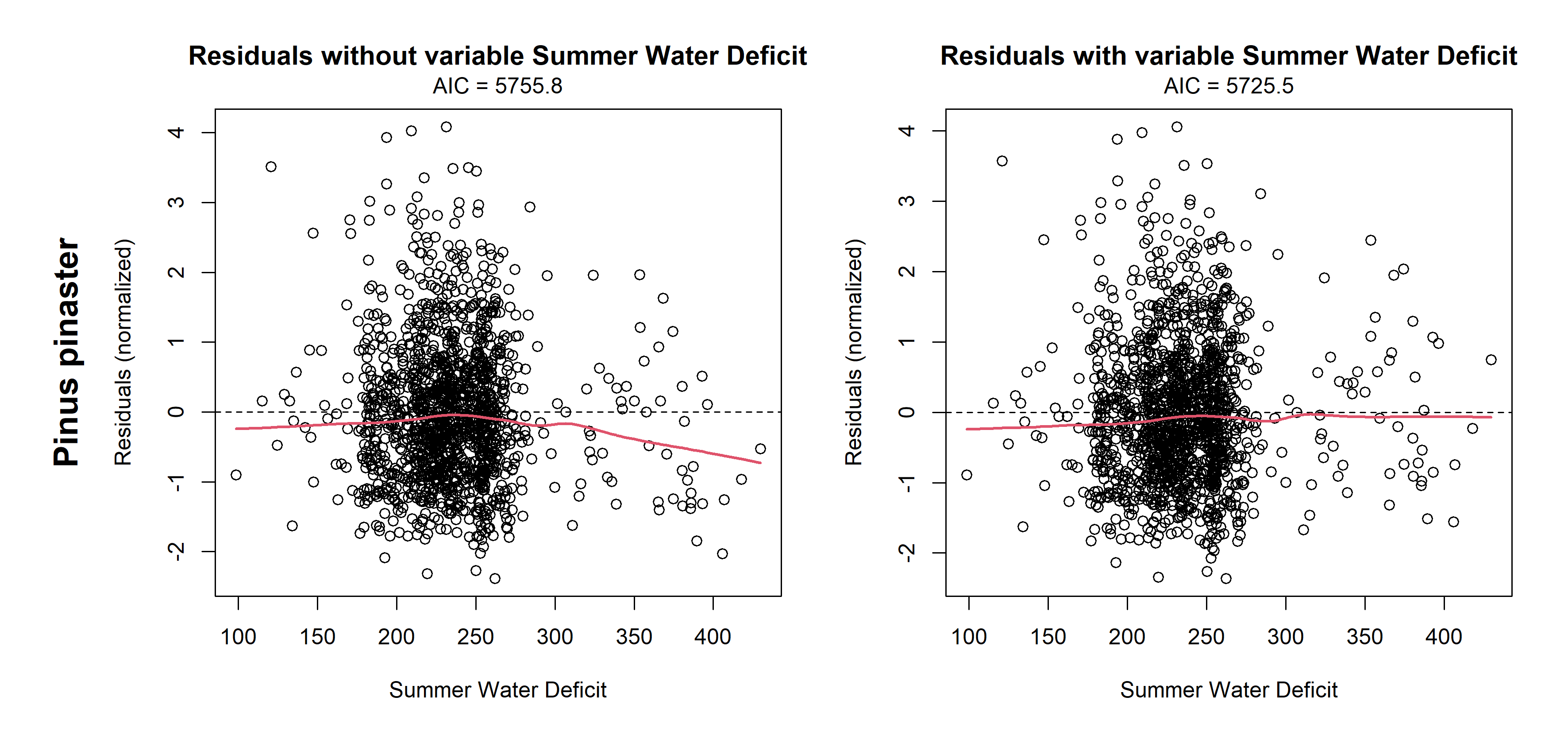

Figure D4. Models normalized residuals against Summer Water Deficit for Pinus pinaster.
On the left side, the residuals are for a model before the inclusion of the Summer Water Deficit variable.
On the right side, the residuals are for the model with the inclusion of the Summer Water Deficit variable.

##### e. Quercus petraea – effect of variable Annual Precipitation

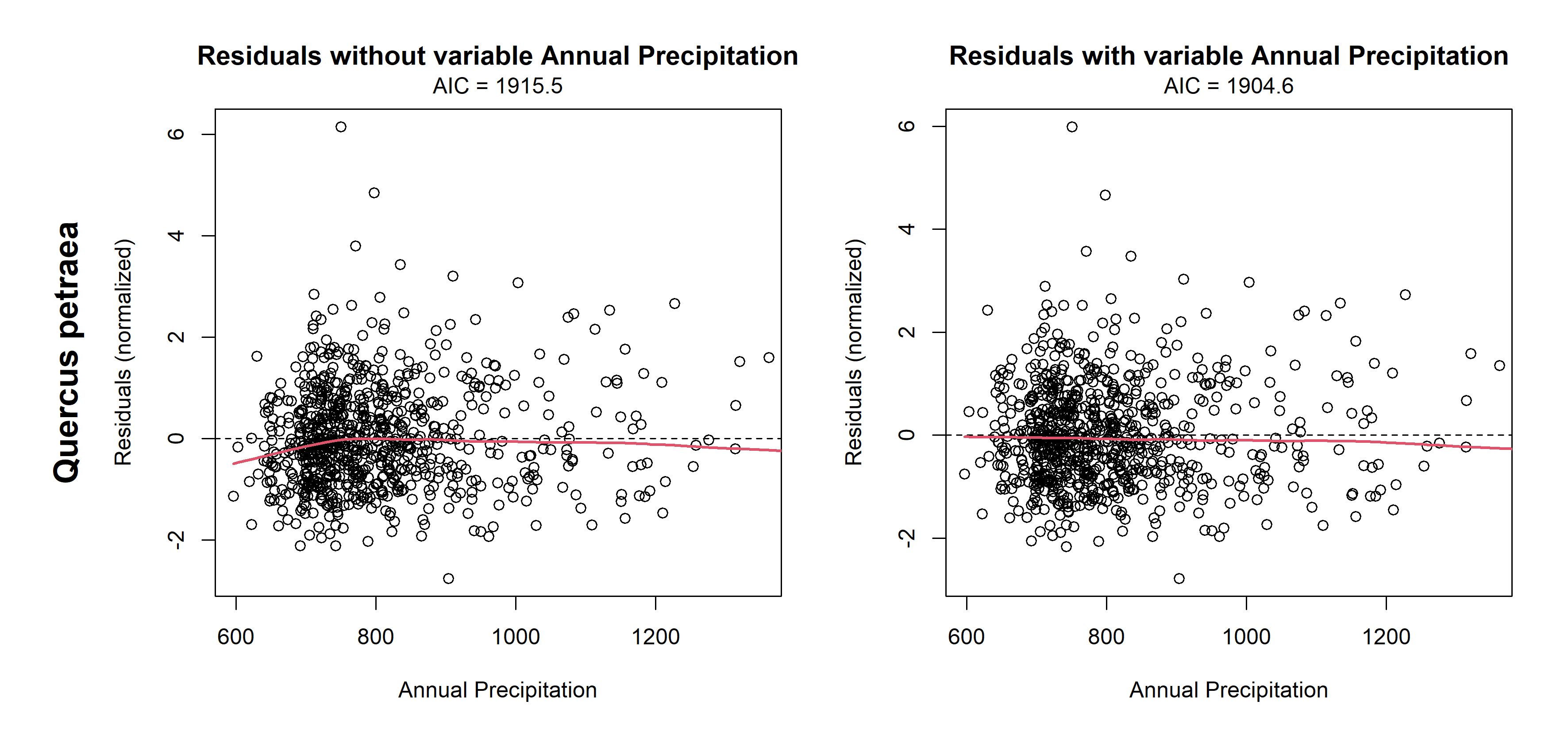

Figure D5. Models normalized residuals against Annual Precipitation for Quercus petraea.
On the left side, the residuals are for a model before the inclusion of the Annual Precipitation variable.
On the right side, the residuals are for the model with the inclusion of the Annual Precipitation variable.

##### f. Quercus petraea – effect of variable Summer Max Temperature

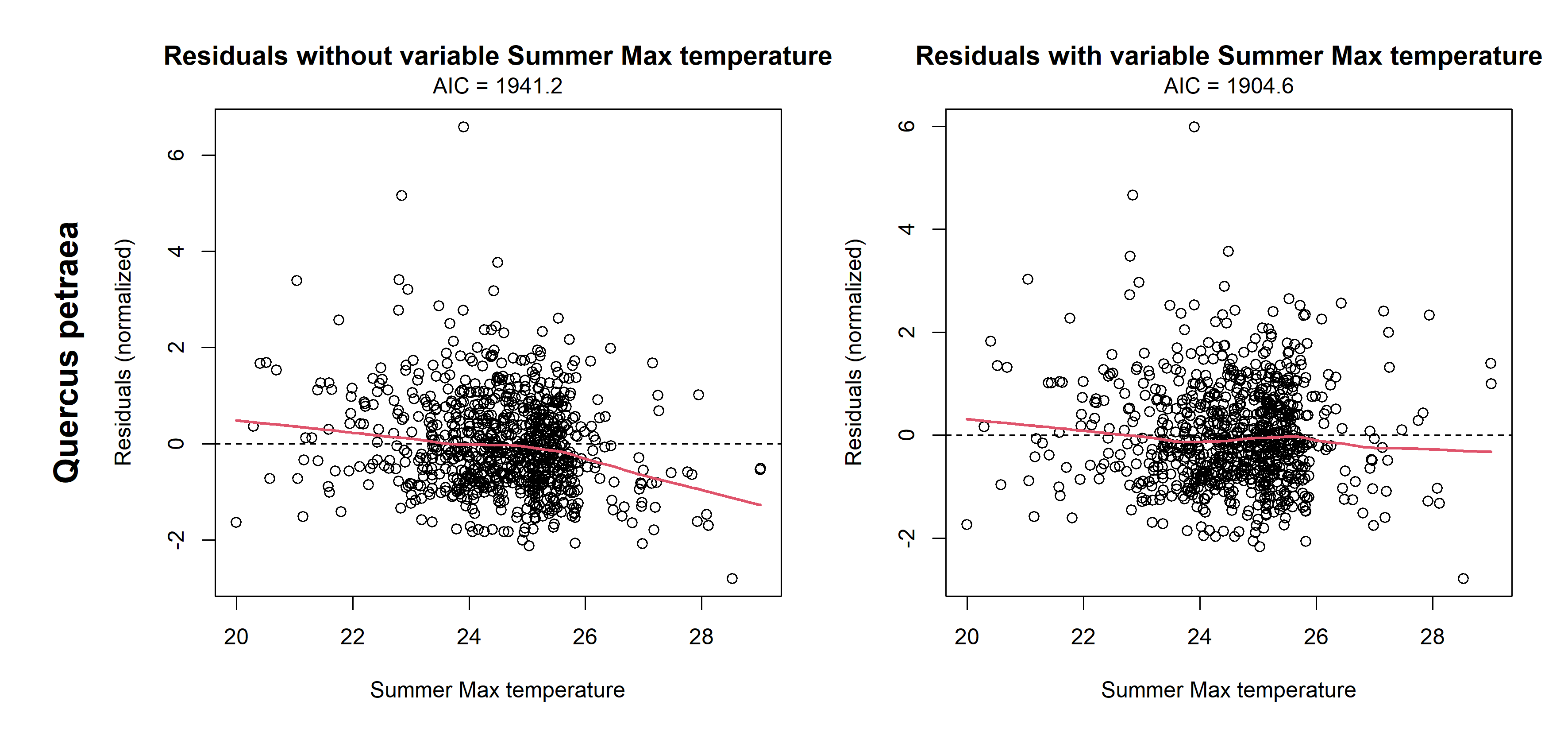

Figure D6. Models normalized residuals against Summer Max Temperature for Quercus petraea.
On the left side, the residuals are for a model before the inclusion of the Summer Max Temperature variable.
On the right side, the residuals are for the model with the inclusion of the Summer Max Temperature variable.

##### g. Quercus robur – effect of variable Summer Water Deficit

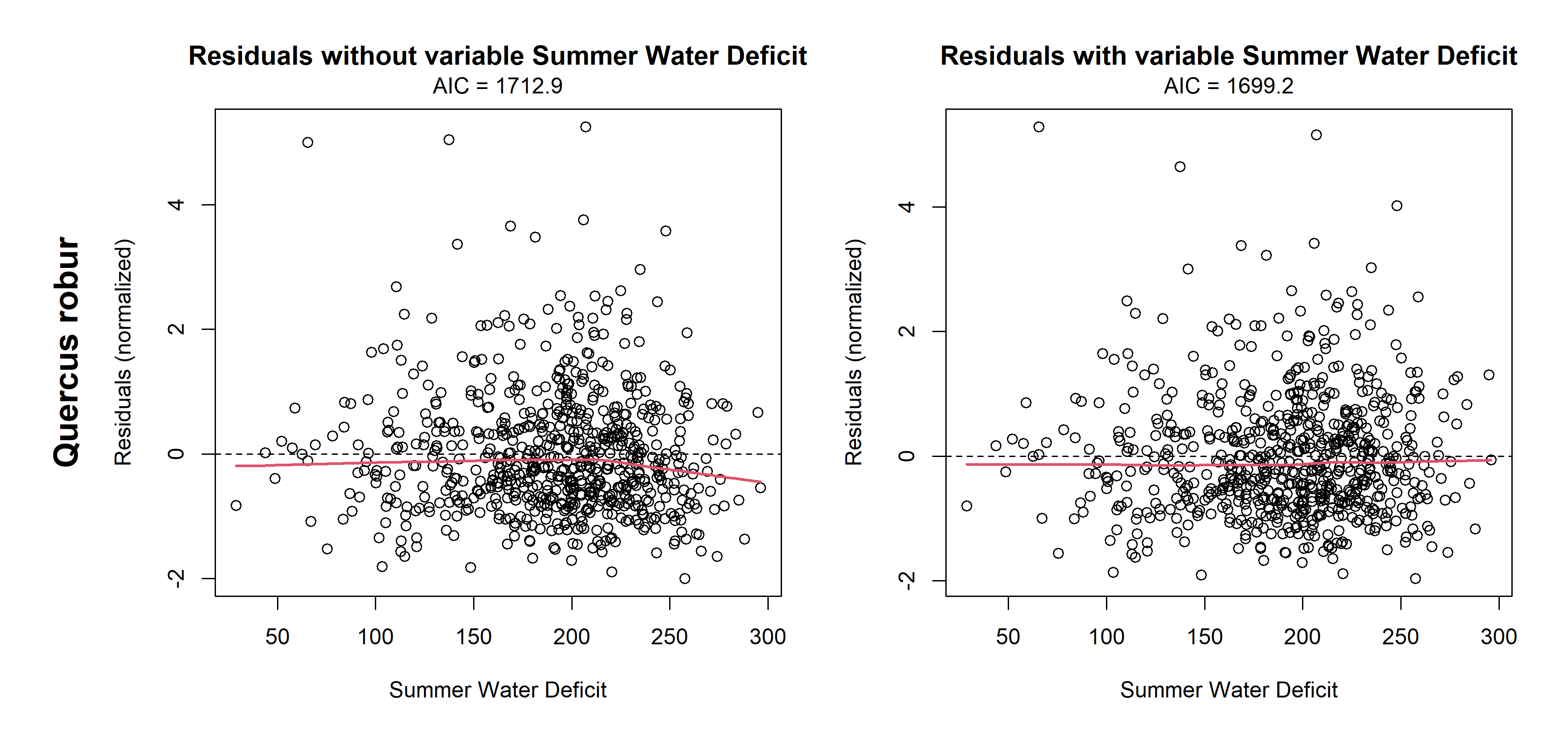

Figure D7. Models normalized residuals against Summer Water Deficit for Quercus robur.
On the left side, the residuals are for a model before the inclusion of the Summer Water Deficit variable.
On the right side, the residuals are for the model with the inclusion of the Summer Water Deficit variable.

1. From flora bioindication (Pinto et al., 2016)
2. <https://inventaire-forestier.ign.fr/?article773>
